# High performance microbial opsins for spatially and temporally precise perturbations of large neuronal networks

**DOI:** 10.1101/2021.04.01.438134

**Authors:** Savitha Sridharan, Marta Gajowa, Mora B. Ogando, Uday Jagadisan, Lamiae Abdeladim, Masato Sadahiro, Hayley Bounds, William D. Hendricks, Ian Tayler, Karthika Gopakumar, Ian Antón Oldenburg, Stephen G. Brohawn, Hillel Adesnik

**Affiliations:** Department of Molecular and Cell Biology, University of California, Berkeley; The Helen Wills Neuroscience Institute

## Abstract

Patterned optogenetic activation of defined neuronal populations in the intact brain can reveal fundamental aspects of the neural codes of perception and behavior. The biophysical properties of existing optogenetic tools, however, constrain the scale, speed, and fidelity of precise optical control. Here we use structure-guided mutagenesis to engineer opsins that exhibit very high potency while retaining fast kinetics. These new opsins enable large-scale, temporally and spatially precise control of population neural activity *in vivo* and *in vitro*. We benchmark these new opsins against existing optogenetics tools with whole-cell electrophysiology and all-optical physiology and provide a detailed biophysical characterization of a diverse family of microbial opsins under two-photon illumination. This establishes a toolkit and a resource for matching the optimal opsin to the goals and constraints of patterned optogenetics experiments. Finally, by combining these new opsins with optimized procedures for cell-specific holographic photo-stimulation, we demonstrate the simultaneous co-activation of several hundred spatially defined neurons with a single hologram, and nearly double that number by temporally interleaving holograms at fast rates. These newly engineered opsins substantially extend the capabilities of patterned illumination optogenetic paradigms for addressing neural circuits and behavior.

## Introduction

Microbial opsins, which flux ionic current in response to illumination, have empowered neuroscientists to causally perturb brain circuits yielding fundamental insight into brain function(Fenno, Yizhar and Deisseroth, 2011). Optogenetics with spatiotemporally patterned illumination enables investigators to causally relate precise features of neural activity with specific aspects of sensation, cognition and action(Ronzitti, Ventalon, *et al*., 2017, Marshel *et al*., 2019, Carrillo-Reid *et al*., 2019, Gill *et al*., 2020, Daie, Svoboda and Druckmann, 2021, Anselmi *et al*., 2011, Dhawale *et al*., 2010, Lutz *et al*., 2008, Fan *et al*., 2020, Packer *et al*., 2012, Russell *et al*., 2019, Zhang *et al*., 2018, Packer *et al*., 2014, Rickgauer, Deisseroth and Tank, 2014, Mardinly *et al*., 2018, Pégard *et al*., 2017, Yang *et al*., 2018, Papagiakoumou *et al*., 2010, dal Maschio *et al*., 2017, Blumhagen *et al*., 2011, Farah, Reutsky and Shoham, 2007, Fenno, Yizhar and Deisseroth, 2011, Vaziri and Emiliani, 2012). However, the biophysical properties of the opsins employed, including their conductance, kinetics, and sensitivity, constrain the type and scale of perturbations that can be made. Although many different opsins have been identified or engineered, writing in precise spatiotemporal patterns of neural activity requires opsins that enable the control of large groups of neurons with high temporal fidelity(Mardinly *et al*., 2018, Marshel *et al*., 2019, Prakash *et al*., 2012, Forli *et al*., 2018). Thus, opsins that are more potent and have kinetic properties enabling high frequency control of spikes trains can substantially expand the capabilities of optogenetic paradigms, particularly in challenging contexts like the mammalian brain. Additionally, since strong over-expression of microbial opsins can alter neuronal morphology(Miyashita *et al*., 2013), achieving high performance neural control with lower levels of opsin expression is desirable. Finally, activating large groups of neurons with multiphoton optogenetics requires high potency opsins due to the much higher energies need to activate opsins with two-photon excitation while avoiding brain heating or tissue damage(Picot *et al*., 2018, Mardinly *et al*., 2018). Furthermore, even one-photon optogenetics applications can benefit owing to thermal constraints during visible light illumination as well(Owen, Liu and Kreitzer, 2019).

Engineering an opsin that provides both potency but also fast closing kinetics is challenging since properties like light sensitivity generally scale inversely with closing kinetics (Mattis *et al*., 2012). For example, the ultrafast ChR2 mutant CheTa(Gunaydin *et al*., 2010) or the ultrafast, red-shifted vfChrimson mutant of Chrimson(Mager *et al*., 2018) exhibit very fast kinetics but have relatively low ionic conductance. Conversely, extremely potent opsins such as the chimeric opsin ReaChR(Lin *et al*., 2013) or the naturally occurring opsin ChRmine(Marshel *et al*., 2019) generate large ionic fluxes but have very slow closing kinetics. ChroME, a point mutant of Chronos(Klapoetke *et al*., 2014, Ronzitti, Conti, *et al*., 2017), somewhat breaks this trend, exhibiting very fast kinetics while still exhibiting high potency(Mardinly *et al*., 2018). Thus, if one could engineer more potent variants of ChroME without substantially slowing its kinetics, such improved opsins will better enable the control of large-scale population activity involved in complex perceptions, cognitive functions and motor actions across species.

We thus set out to engineer new opsins based on the ChroME backbone with significantly enhanced potency while retaining fast kinetics. We then benchmarked these new variants against existing tools with a broad set of electrophysiological and optical approaches. In particular, we present two new mutants (‘ChroME2f’ and ‘ChroME2s’) with enhanced properties that can support large scale, temporally precise multiphoton excitation in the intact brains of awake animals. These enhanced opsins provide up to a 5-7-fold increase the number of simultaneously controllable neurons compared to ChroME, yet still provide sub-millisecond temporal control at high firing rates.

All-optical perturbation experiments, where the experimenter both controls and measures the activity of neurons with light, is emerging as a powerful tool in neuroscience. However, a main challenge for these experiments is the possibility that the laser used for two-photon imaging of neural activity (e.g., via GCaMP sensors) might incidentally depolarize the neurons of interest through activation of the opsin molecules(Packer *et al*., 2014, Mardinly *et al*., 2018, Forli *et al*., 2018), which we refer to as ‘optical cross-talk’. Since opsins absorb at the typical wavelength for GCaMP imaging (∼920-930 nm, see Fig. 6 below), this is an important concern for any all-optical experiment using this or other green sensors. We therefore also carefully quantify unwanted ‘optical cross-talk’ across a broad range of conditions to define the limits under which functional imaging can be conducted while generating minimal unwanted depolarization. Furthermore, we provide a comprehensive spectral survey of opsins under two-photon illumination across a wide band (750-1300 nm) that can help experimenters choose among these potent opsins depending on the imaging wavelength to minimize or reduce spectral overlap with the activity sensor. These data provide an essential knowledge base for the further refinement of these opsins and their future use with the ever-expanding toolbox of activity indicators for specific ions, voltage, and neurotransmitters. Finally, by leveraging ChroME2.0 we demonstrate the simultaneous optogenetic perturbation of 300-400 of unique cortical pyramidal neurons at a time, and >600 unique neurons per second in the brains of awake mice with multiphoton holographic optogenetics.

## Results

### Structure-guided design of ultra-potent, ultra-fast opsins based on ChroME

To engineer an ultrapotent opsin that retains fast kinetics we exploited the opsin ChroME, which is a point mutant of the ultra-fast opsin Chronos. Using ChroME as a backbone we targeted several key residues based on structural homology modeling that are likely to either line the channel pore and/or interact with the retinal chromophore, and thus could strongly influence channel biophysics(Kato and Nureki, 2013, Kato *et al*., 2012) (Fig. 1a). We generated 9 single point mutants, and first characterized each one in cultured CHO cells with visible light illumination to rapidly assay their estimated potency, kinetics, and spectral properties (Fig. 1bw-1d, 1i).

**Figure 1.**
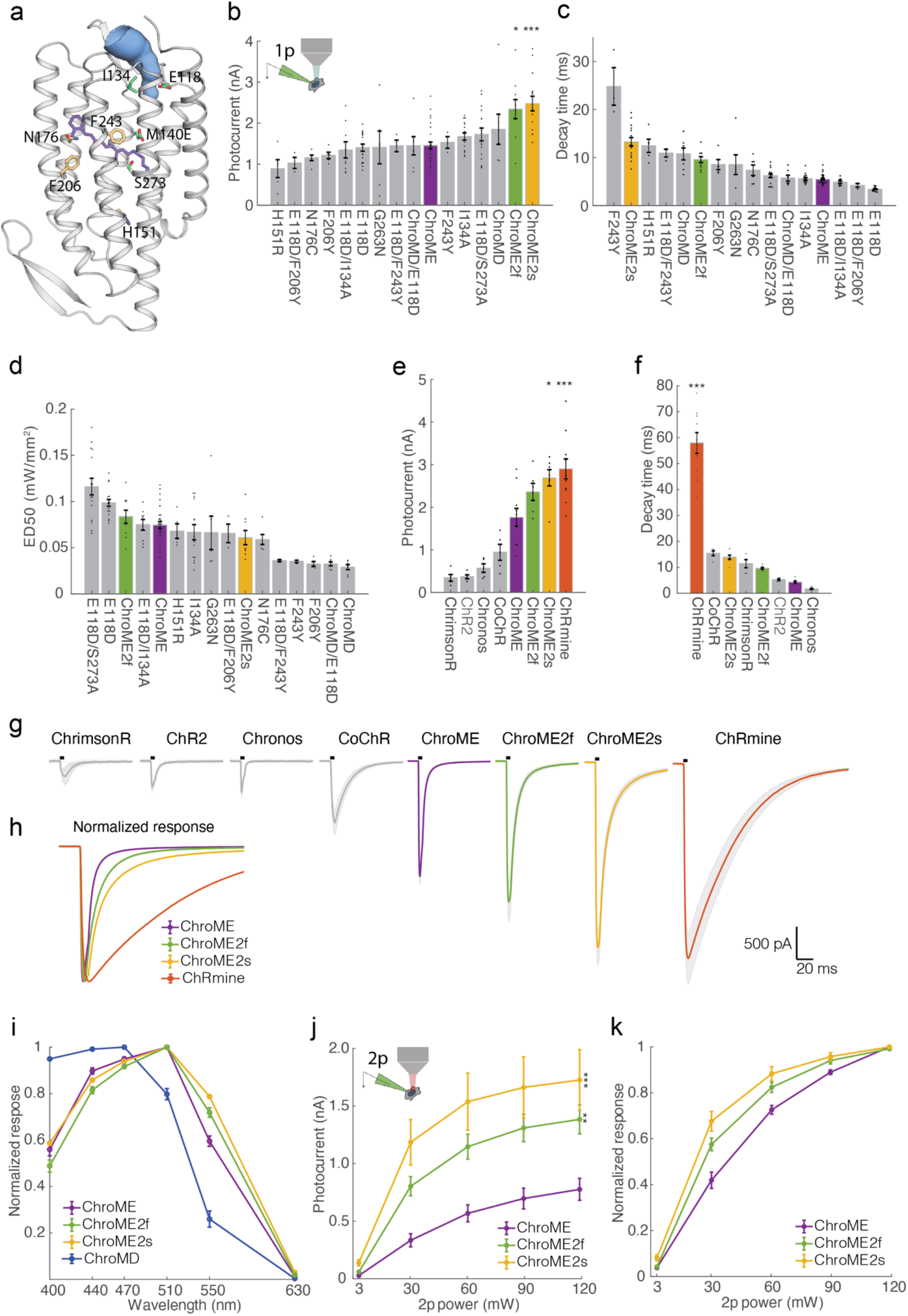
Design and characterization of second generation ChroME variants under visible and two-photon excitation. **a)** Model of ChroME showing some of the amino acids that targeted for mutagenesis. The retinal chromophore is shown in purple and the putative ion channel pore is indicated in blue. **b)** Inset: schematic of a transfected cultured Chinese hamster ovary (CHO) cell under whole cell patch clamp. Peak photocurrents recorded under for each opsin mutant at (power = 0.45 mW at 510 nm unless otherwise indicated). For ChroMD, ChR2 and CoChR, 470 nm was used and for ChrimsonR, 630nm was used). **c)** Decay time and **d)** Sensitivity (ED50) of opsin mutants from b) **e)** Peak photocurrents of ChroME2f and ChroME2f compared against a panel or previously characterized opsins. **f)** Estimated decay times from monoexponential fits for the same panel of opsins as in e). **g)** Average photocurrents traces for indicated opsins obtained from data shown in e). The black bar above the trace indicates a 5 ms pulse of light. **h)** Peak-normalized traces of the indicated opsins obtained from data shown in e. **i)** Visible wavelength spectra of the indicated opsins at the specified wavelengths. **j)** Photocurrents at the indicated powers under two-photon illumination at 1040 nm for the indicated opsins. **k)** Normalized two-photon photocurrent response for the photocurrents show in j). Data are presented as mean and s.e.m. Statistics: *P < 0.05, and ***P < 0.001 with ChroME as reference; 1-way ANOVA with multiple comparisons test.

Many mutations of ChroME diminished the effective photo-conductance and/or slowed channel properties. However, several mutants substantially either sped up channel closing or increased whole-cell photocurrents which are both desirable characteristics. Three mutants had notable qualities: a glutamate to aspartate mutation at site 118 in ChroME substantially sped up the kinetics (decay time ChroME: 5.48 ms, ChroME E118D 3.42 ms, Fig. 1c), while an isoleucine to alanine at 134 increased peak photocurrents (ChroME: 1.46 nA; ChroME I134A: 1.67 nA, Fig. 1b). A serine to alanine mutation at site 273 likewise significantly potentiated whole-cell photocurrents (ChroME: 1.46 nA; ChroME S273A: 2.48 nA, Fig. 1b) but at the expense of a slowing of channel kinetics (decay time ChroME: 5.48 ms, ChroME S273A 13.28 ms, Fig. 1c). We refer to ChroME point S273A as ‘ChroME2s’ since it exhibited the highest potency of all the mutants we tested as well as improved sensitivity (Fig. 1d), but with somewhat slower kinetics.

We therefore hypothesized that combining these mutations could yield an opsin that is substantially more potent than ChroME while retaining very fast kinetics. Indeed, we found that the ChroME triple mutant S273A, E118D, I134A had substantially stronger currents (ChroME: 1.46 nA; ChroME S273A, E118D, I134A: 2.34 pA, 1b) and partially rescued the decay time increased by the S273A mutation (ChroME-S273A: 13.28 ms; ChroME S273A, E118D, I134A: 9.58 ms, 1c). We hereafter refer to this opsin as ‘ChroME2f’ (‘f’, for ‘fast’). Finally, we identified several other mutants with unique and potentially advantageous properties, including a variant termed ‘ChroMD’ (Chronos with a methionine to aspartate mutation at site 140) that exhibited a substantial blue-shift in its absorbance peak (Fig. 1i) as well as a strong increase in light sensitivity (Fig 1d). We chose to focus our efforts on ChroME2f and ChroME2s owing to their high potency, overall optimal kinetics and similar absorbance spectra with ChroME (Fig 1i) under visible illumination.

We then sought to compare these new ChroME variants with the standard opsins that are currently in use such as Chronos, ChR2, ChrimsonR, CoChR and ChRmine under one-photon excitation in cell culture. The ChroME variants, ChroME2f and ChroME2s, were among the most potent opsins with respect to photocurrents in our comparison (Fig. 1e). ChRmine, which provides greater photocurrents than ChroME(Marshel *et al*., 2019) (Fig. 1e), was slightly stronger than even the ChroME2f/s variants (Fig. 1e). However, the closing kinetics of ChRmine were much slower: about 6-fold slower than ChroME2f and 4-fold slower than ChroME2s (Fig. 1f-h). The ChroME2f/s variants also outperformed ChroME under two-photon excitation (Fig. 1j-k). To test how ChroME2f and ChroME2s would perform in neurons, we next compared photocurrents and light-evoked spiking in L2/3 cortical neurons expressing ChroME, ChroME2s, ChroME2f. We expressed the opsins in mouse L2/3 cortical pyramidal neurons by *in utero* electroporation and activated them with both one-photon (Fig. 2a) and two-photon excitation (Fig. 2b, c). Whole-cell recordings confirmed that ChroME2s and ChroME2f both provided substantially larger photocurrents than ChroME (Fig. 2a, b) and closely recapitulated decay times that were observed in our original screen by cell culture (Fig. 2c).

**Figure 2.**
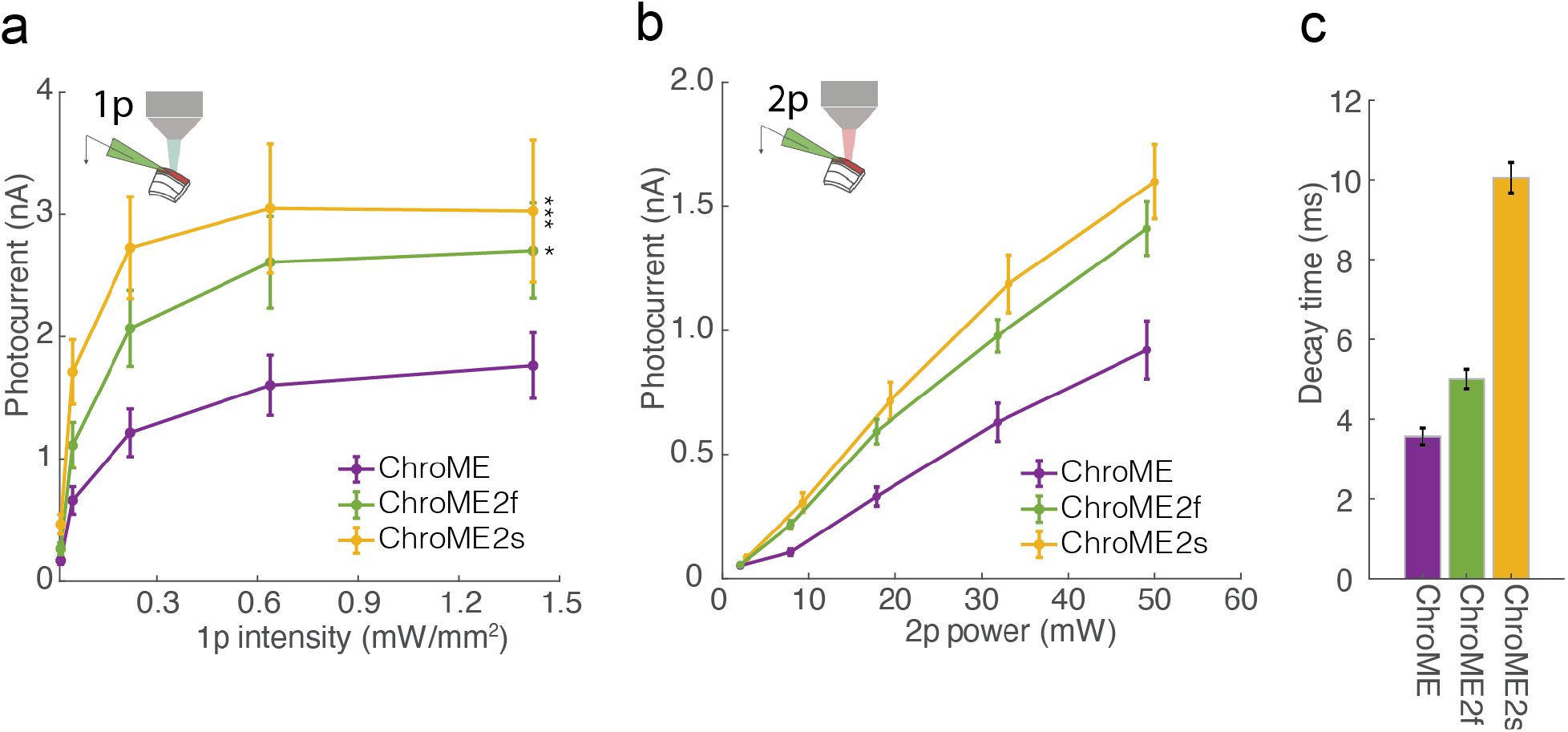
Validation of ChroME variants in acute brain slices under visible and two-photon excitation. **a)** Photocurrents under full-field 1p illumination at 510 nm and the indicated powers from L2/3 pyramidal neurons expressing the indicated opsins via in utero electroporation. P < 0.05, and ***P < 0.001 with ChroME as reference; 1-way ANOVA with multiple comparisons test. **b)** As in a) but with two-photon excitation and a ∼12.5 mm diameter spot. **c)** Decay times of the three opsins measured with monoexponential fits (P<0.05, One-way ANOVA, all groups significantly different except ChroME and ChroME2f with multiple comparisons correction). Data are presented as mean and s.e.m. Statistics: *

### ChroME2.0 variants provide high potency, high temporal fidelity control over neural activity

Based on these results, we next compared the light powers needed to reliably drive neurons to action potential threshold across a range of specific frequencies. We activated L2/3 pyramidal neurons with pulse trains (pulse width = 5 ms) of varying frequency and power, as well as compared the electrophysiological response to single pulses of increasing power and duration (Fig. 3). For these experiments we also included ChRmine since it’s among the most potent opsins identified yet(Marshel *et al*., 2019). Although all four opsins could reliably drive all recorded neurons to fire, we observed substantial differences in the power and frequency response of cortical neurons depending on the opsin used. Both ChroME2f and ChroME2s could drive neurons to spike at substantially lower powers than could ChroME, yet readily maintained high frequency firing up to the highest frequency tested (40 Hz, Fig. 3b,c,e). Particularly for ChroME2f, the ability of light pulses to reliably generate spikes was remarkably robust to changes in pulse frequency (5-40 Hz) at each light power tested (Fig. 3e). Importantly, for all ChroME-based opsins light pulses generated one spike per pulse under most conditions (Fig. 3e). ChRmine, in contrast, could evoke spiking at even lower power levels than the ChroME2 variants, but the spiking response was not a monotonic function of power and response levels were less reliable across frequencies (Fig. 3d, e). Since a prior study employed shorter pulses to activate ChRmine-expressing neurons(Marshel *et al*., 2019), we collected an additional data set with pulse trains set using 0.5 ms pulses. Under these conditions the dynamic range of activating ChRmine-expressing neurons appeared larger with respect to laser power, and the response function was monotonic with power (Figure S1a, n = 5). However, more power per neuron was needed to generate these spikes than when using 5 ms pulses, and at low frequencies (5 Hz) high power 0.5 ms pulses still generated more than one spike (Figure S1a).

**Figure 3:**
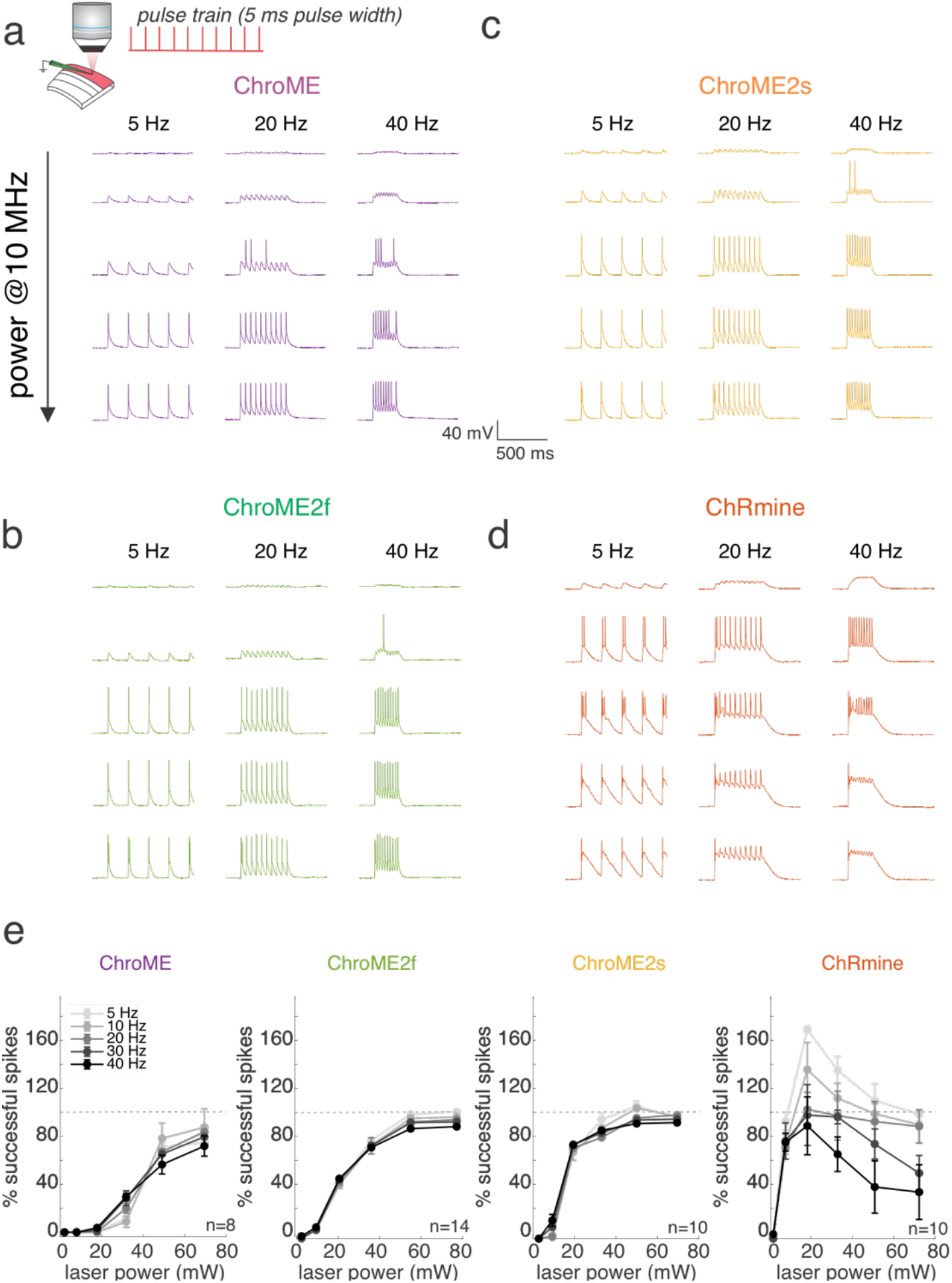
Comparison of two-photon light-evoked spiking of L2/3 pyramidal neurons expressing ChroME2.0 variants to ChroME and ChRmine. **a)** Top: Schematic the experiment. A single L2/3 neuron is patched in current clamp mode and illuminated with fixed frequency trains of 5 ms pulses at 1040 nm, 10 MHz repetition rate. Bottom: example traces from a ChroME-expressing neuron, **b-d)** Example traces from representative ChroME2f, ChroME2s, and ChRmine-expressing cells. **e)** Plots of the fraction of 5 ms light pulses that drove spikes across laser powers and stimulation frequencies for the four opsins. Sample size (cells) is indicated in the panels. The two-photon excitation spot size was 12.5 mm in diameter. Data points represents the mean +/-s.e.m.

Finally, we compared how each opsin could drive cells as a function of illumination time for single pulses ranging between 1-30 ms (Figure S1b-e). For all opsins, increasing the duration substantially increased spike probability of expressing neurons as a function of power.

To obtain a better understanding of how these different opsins would influence the temporal fidelity of light induced spike trains in more physiological conditions, we stimulated opsin-expressing neurons with broadband (‘poisson’) trains of light pulses (pulse width = 5 ms) while injecting noisy sub-threshold currents to mimic the synaptic bombardment neurons experience under *in vivo* conditions (Fig. 4). This approach allowed us to quantify how spike latency, jitter, and the probability of spike success varied with instantaneous spike frequency. The laser power for each neuron was set above the level needed to reliably evoke an action potential with a single isolated 5 ms pulse to control for variation in opsin expression and intrinsic excitability across neurons. We found that ChroME2f or ChroME2s-expressing neurons could reliably follow pulse trains with sub-millisecond jitter across a wide range of frequencies. (Figure 4a-f), while ChRmine-expressing neurons dropped to about 50% spike success even at low frequencies (Fig. 4f). We retested ChRmine-expressing neurons with a pulse width of 0.5 ms, but under these conditions the neurons still did not reliably follow the random, broadband pulse trains (Figure S2). These data demonstrate that ChroME2f and ChroME2s can enable the precise reproduction of physiological-like spike trains across a broad range of frequencies.

**Figure 4:**
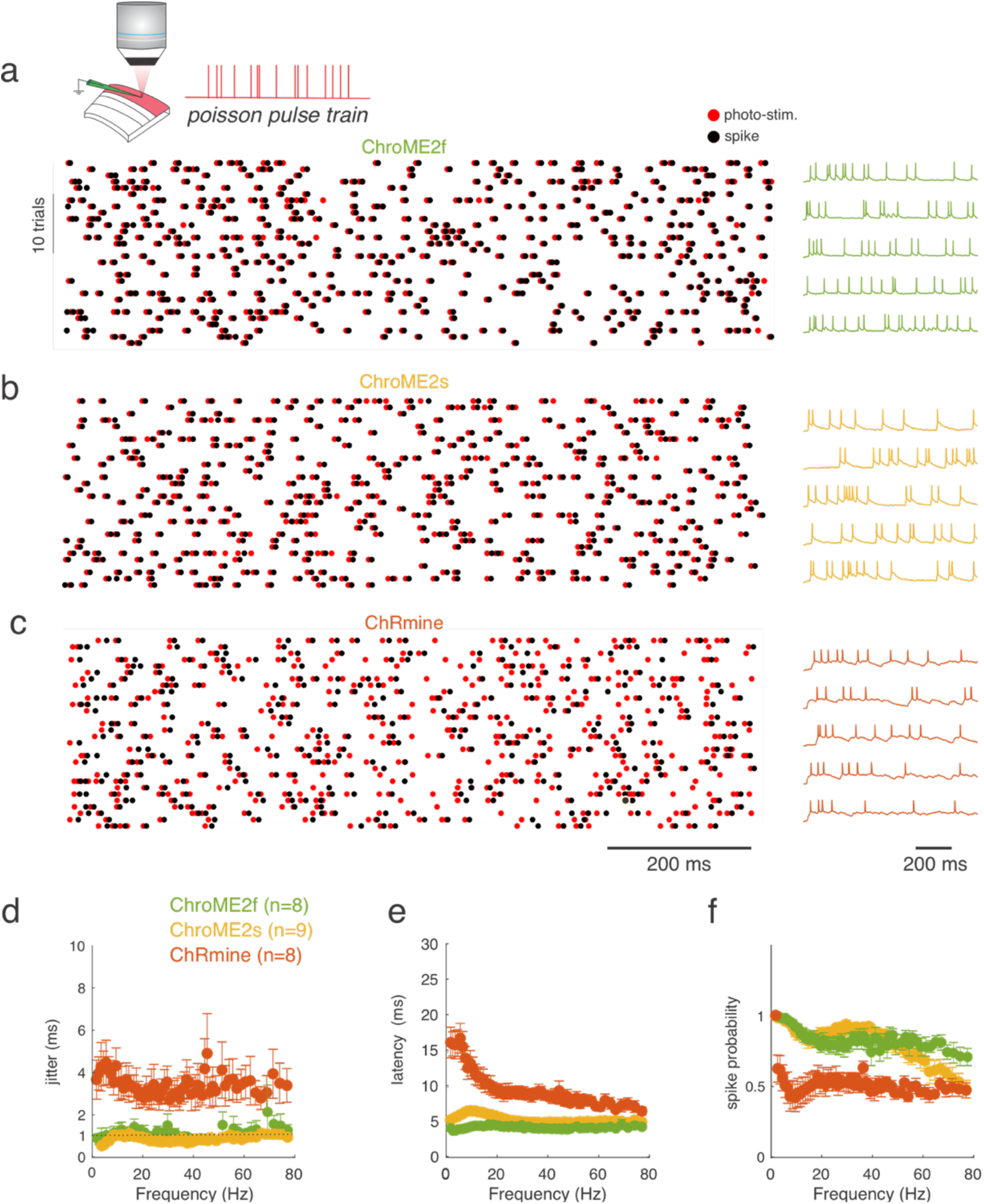
Temporal fidelity of light evoked spiking with broadband ‘poisson’ stimulus trains. **A)** Top: Schematic the experiment. A single L2/3 neuron is patched in current clamp mode and illuminated with poisson-like trains of 5 ms pulses at 1040 nm, 10 MHz repetition rate. Bottom: example raster plot of light pulses (red) and spikes (black) from a ChroME2f-expressing neurons. Right: five example traces from the recording. **B**,**C)** Example spike rasters (left) and traces from neurons expressing ChroME2s or ChRmine. **D-F)** Plot of the jitter, latency and spike probability of light-evoked spikes during the broadband stimulation across the three indicated opsins as a function of instantaneous pulse frequency. Sample size is indicated in the panel. Data represent the mean +/-s.e.m.

### Optical cross-talk and spectral characteristics of the new and existing opsins

The spectral absorbance of opsins delineates the optimal optical wavelength to use for their photo-excitation, but also critically determines how strongly they are incidentally activated by a second two-photon imaging laser. Few opsins have been fully spectrally characterized in the two-photon regime since most prior studies have employed Ti:Sapphire lasers that have a limited tuning range. Thus, we acquired absorbance spectra (820-1300 nm, Insight X3) for a large panel of opsins with diverse biophysical features to provide a critical knowledge base for the future selection of opsins for two-photon optogenetics experiments. We expressed 13 different opsins in cell culture and illuminated the cells with a soma-sized spot of light carefully calibrated for the intrinsic power spectrum of the laser source and transmission of the microscope (Fig. 5). ChroME, ChroME2f, and ChroME2s all exhibited fairly similar two-photon excitation spectra with a peak of ∼1000 nm, substantial absorbance at 920 nm, and minimal absorbance beyond 1200 nm. We characterized other well-known opsins including ChRmine, ChrimsonR, Chronos, CoChR, ReaChR and ChR2 that have also been used for two-photon excitation (Fig. 5). In general, their two-photon excitation matched well with partial spectra taken in prior work where available. The blue-shifted ChroME mutant, ChroMD, showed a strong blue shift consistent with the 1p absorbance data described above. Additionally, we characterized the more recently identified PsChR, which is perhaps the most blue-shifted cation opsin yet described. Finally, we obtained complete spectra for the potent anion opsins GtACR1 and GtACR2. These data establish a palette of opsins for diverse experiments and help outline possible multi-spectral for bidirectional optogenetics.

**Figure 5:**
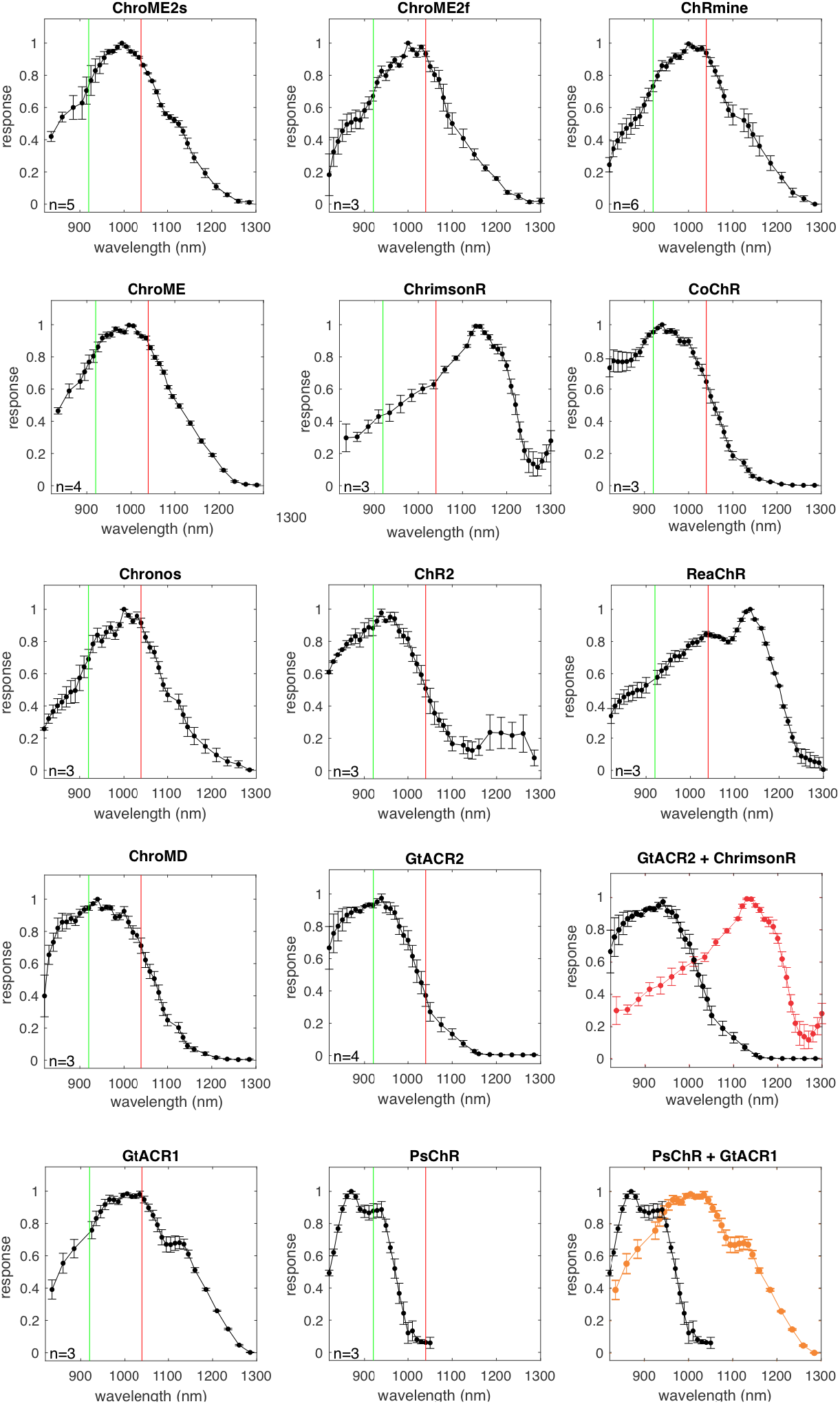
Two-photon excitation spectra of various opsins and ChroME2.0 variants. For each opsin the plot shows the peak-normalized photocurrent recorded in CHO cells transfected with the indicated opsin using an Insight X3 tunable laser (80 MHz). The green vertical line marks 920 nm, while the red line indicates 1040 nm.

**Figure 6:**
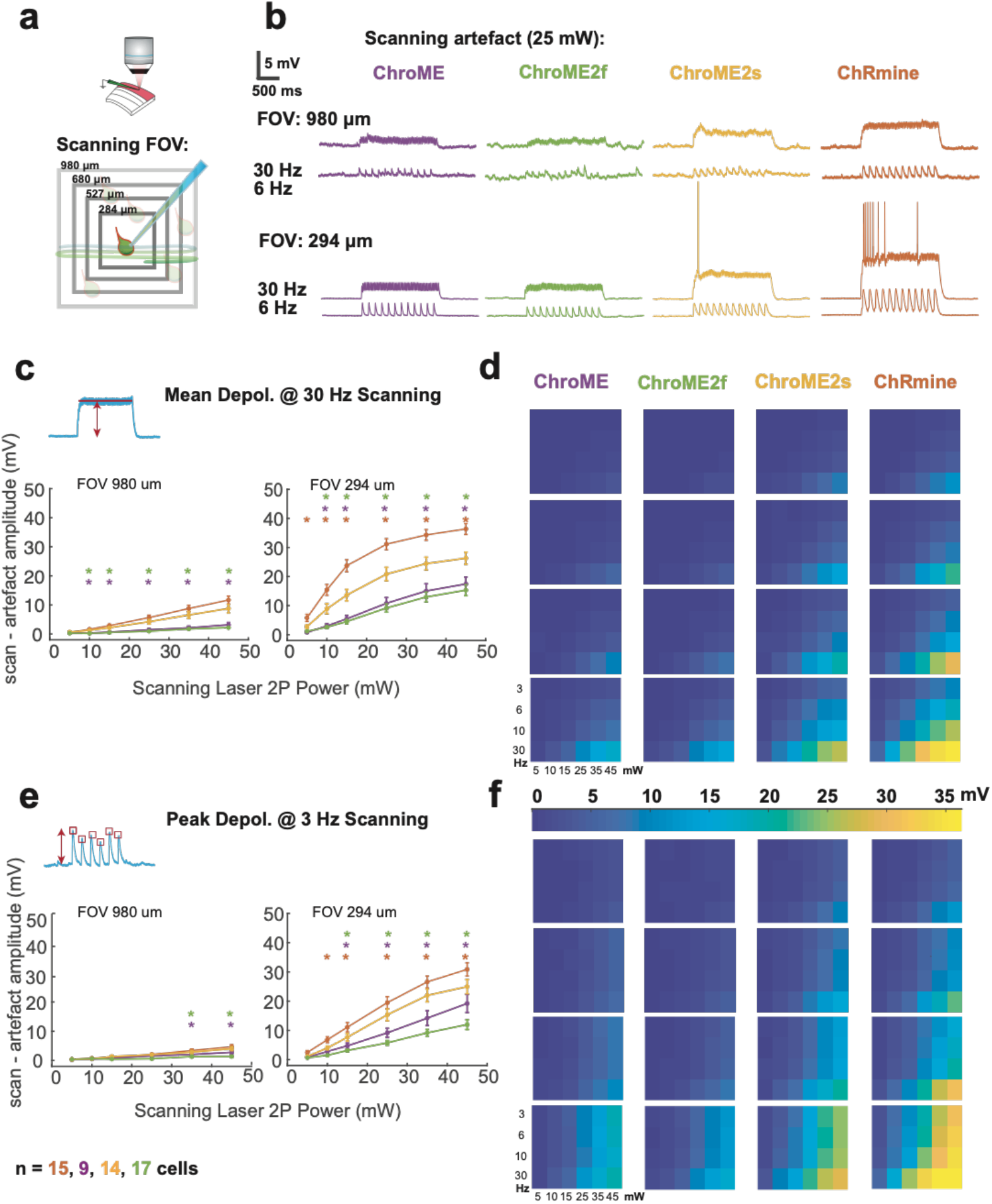
Activation of opsin variants by resonant galvo scanning. **A)** Schematic of the experiment indicating a patched neuron expressing opsin and raster scanning the imaging laser at different optical zooms. **B)** Example membrane potential traces from four neurons expressing ChroME, ChroME2f, ChroME2s, and ChRmine at two different zoom levels and two different imaging rates. Laser power = 25 mW, 920 nm. **C)** Plot of the mean scanning artifact amplitude across a sample of neurons for each of the four opsins for two different zoom levels and six different laser powers at 3Hz frame rate. **D)** Heat map plot for scanning artefact across all 96 conditions. **E**,**F)** Same as for C,D but for the peak depolarization computed across the entire the trial. Sample size per opsin indicated ta bottom. Error bars are s.e.m. Sample size is indicated in the panel.

Next, to quantify optical cross-talk that could occur during all-optical experiments, we made whole-cell current-clamp recordings of cortical neurons expressing each opsin, while imaging the slice with a resonant-scanning galvo/galvo system (Fig. 6a). We systematically varied three key determinants of optical cross-talk: imaging power, field of view, and frame rate. We found that under imaging conditions that are commonly used to sample cortical neurons volumetrically (FOV = 980 x 980 microns, frame rate = 6 Hz) neurons expressing ChroME or ChroME2f showed minimal scan-induced depolarization (peak = 1.1+/-0.26 mV, mean: 0.27+/-0.08 mV @25 mW for ChroME, peak = 0.73+/-0.08 mV mean: 0.29 +/-0.05 mV @25 mW for ChroME2f) (Fig. 6b, d,f, Figure S3,S4). Neurons expressing ChroME2s and ChRmine showed somewhat larger depolarization (ChroME2s: peak = 2 +/-0.33 mV, mean: 1.05 +/-0.2 mV @25 mW, ChRmine: peak = 2.4 +/-0.4 mV, mean: 1.1 +/-0.2 mV @25 mW, one-way analysis of variance (ANOVA) p=0.0002, Fisher’s Least Significant Difference tests used for all multiple comparisons, Fig. 6b, d,f, Fig. S3,4).

In smaller FOVs at higher magnifications (294 x 294 microns) which are often used for imaging sub-cellular structures such as dendrites, spines and axonal boutons, depolarizations caused by the scanning beam were substantially larger (Fig. 5b-f). In many ChRmine-expressing and some ChroME2s-expressing neurons the scanning directly evoked spiking from resting potential, but this did not occur in neurons expressing ChroME or ChroME2f (Figure S5). We also measured the intrinsic physiological properties of the neurons under study but did not find substantial differences (Fig. S6). These data provide critical details that can guide which of these opsins to choose for distinct classes of all-optical multiphoton optogenetics experiments and reflect the tradeoffs between opsin potency and kinetics and optical cross-talk during two photon imaging.

### In vivo performance and large-scale control of neural activity with ChroME2.0 variants

In vivo performance and large-scale control of neural activity with ChroME2.0 variants Next, we sought to compare the performance of the most potent opsins identified above for in vivo holographic two-photon optogenetic control. We expressed ChroME, ChroME2s, ChroME2f, ChRmine and ChrimsonR in L2/3 pyramidal neurons via AAV viral transfection (using the template vector AAV-CAG-DIO-[Opsin]-ST-P2A-H2B-mRuby3) in mice expressing GCaMP6s transgenically in cortical excitatory neurons. We found that their potency measured in brain slices was largely consistent with their potency for photo-stimulation in vivo (Fig. 7). ChroME2s and ChroME2f-expressing neurons were both substantially more sensitive than ChroME: at 25 mW/cell the fraction of neurons that could be photo-activated increased ∼5-fold for ChroME2f and ∼7-fold for ChroME2s; at 50 mW/cell, the increase was 2-3 fold compared to ChroME (Fig. 7 c,d) ChRmine exhibited the highest potency among this panel, slightly outperforming ChroME2s, while ChrimsonR exhibited the weakest potency (fraction photoactivable at 50mW, ChrimsonR: 0.059 ± 0.015, ChroME: 0.33 ± 0.025, ChroME2f: 0.64 ± 0.055, ChroME2s: 0.83± 0.004, Chrmine: 0.93± 0.011, p<0.001, one-way ANOVA, p<0.05 for all comparisons, post hoc multiple t-test with holm correction, Fig. 7c, Fig. S7).

**Fig. 7:**
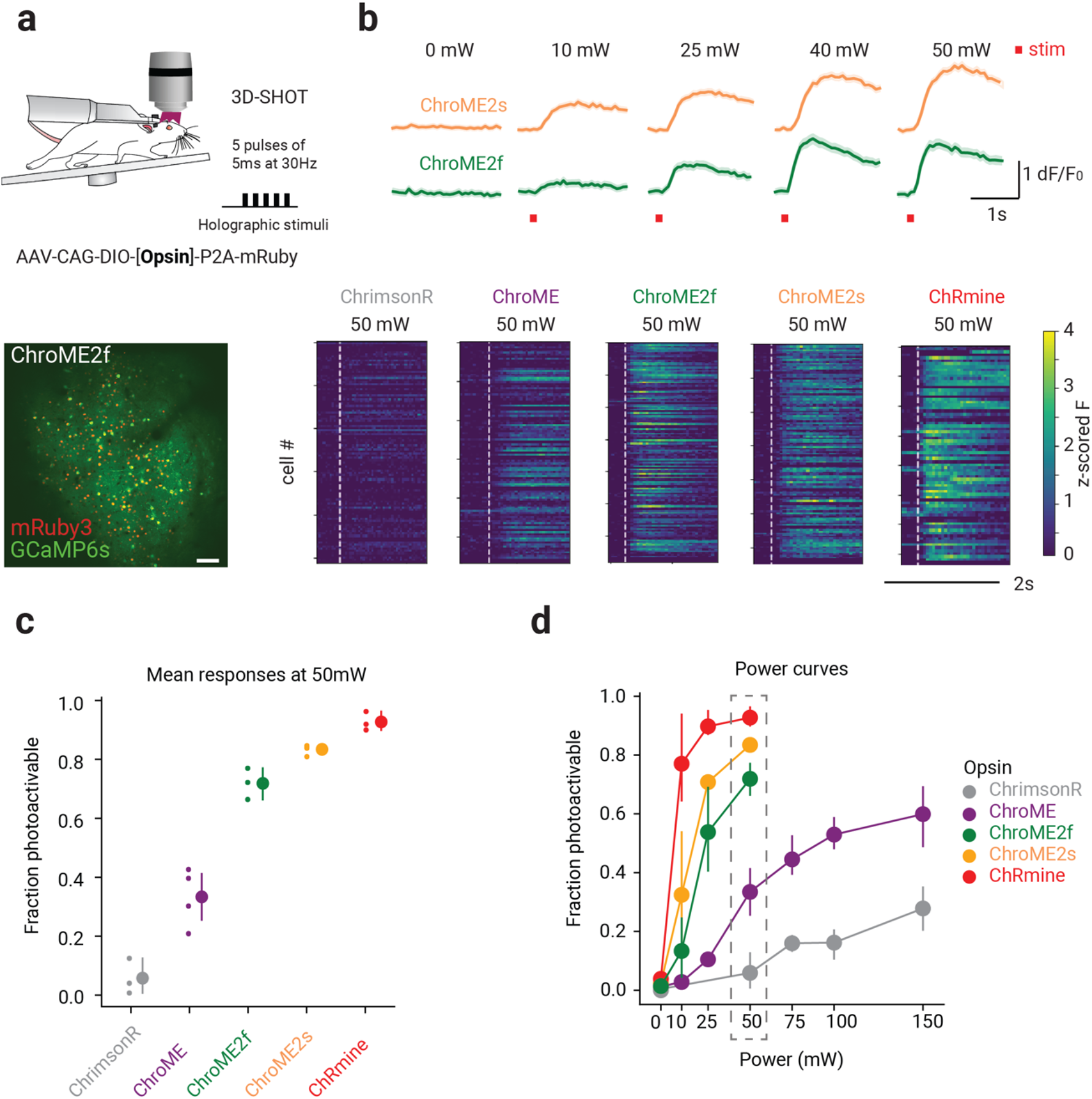
*In vivo* all-optical comparison of opsin potency for photo-stimulation. **A)** Top: Experimental schematic. Adult mice virally expressing each of the opsins in excitatory cells were holographically stimulated using 3D-SHOT. Individual cells were excited with a train of 5 pulses of 5ms duration at 30Hz. Bottom: *In vivo* two-photon image of a representative FOV with all excitatory neurons expressing GCaMP6s (green). Opsin-expressing neurons are labeled with nuclear mRuby3 (red). Scale bar: 100mm. **B)** Top: Example traces of mean responses of cells expressing ChroME2s (Top, Yellow) and ChroME2f (Bottom, green) to different power intensities. Bottom: Representative mean z-scored fluorescence peristimulus time histograms (PSTHs) for opsin expressing cells stimulated at 50mW. Dashed lines indicate the onset of photo-stimulation. **C)** Mean fraction of photoactivable cells (relative to all targeted cells in a FOV) for each opsin at 50mW (***p<0.001, 1-way ANOVA, posthoc multiple T-test with holm correction: Chrimson<ChroME<ChroME2f<ChroME2s<ChRmine, *p<0.05 for all comparisons, N ≥ 3 mice per opsin, with ≥50 cells per mice). **D)** Fraction of photoactivable cells as a function of power for each opsin (***p<0.001, 2-way ANOVA).

Finally, we sought to determine an upper bound on how many neurons we could simultaneously co-activate in a large volume *in vivo* under these expression conditions (Fig. 8) and with a maximal total instantaneous laser energy at the sample of ∼4W. Since ChroME2s is nearly as potent as ChRmine, but still provides excellent control over high frequency firing, we focused on this opsin and designed an optogenetic paradigm to maximize the number of neurons that could be simultaneously photo-stimulated at a single moment (with a single phase mask on the spatial light modulator, SLM) or to maximize the number of neurons that could be co-activated within a defined time window (e.g, one second) by rapidly interleaving multiple SLM phase masks (Fig. 8b). We systematically varied pulse duration, frequency, and power to identify the conditions which would optimize the size of the activated ensemble under both of these conditions (Fig. S8). To further optimize this, we selected opsin-expressing neurons that showed higher light sensitivity. To this end, we systematically probed the optogenetic gain of >1,000 neurons in the brain volume, selected the top 30% of excitable neurons, and then adjusted the power directed to each neuron in a multi-spot hologram to drive a significant increase in calcium signal (Fig. 8d, Figure S8c). Figure 8c summarizes the hologram sizes used across experiments and the resulting photoactivation rates. In one example animal under these conditions we could simultaneously increase the activity of 335 neurons for a 408-target hologram (Fig. 8d,e; Wilcoxon rank-sum test, one-tailed p<0.025). The remaining 73 neurons either did not show a detectable change in calcium signal or were suppressed.

**Figure 8:**
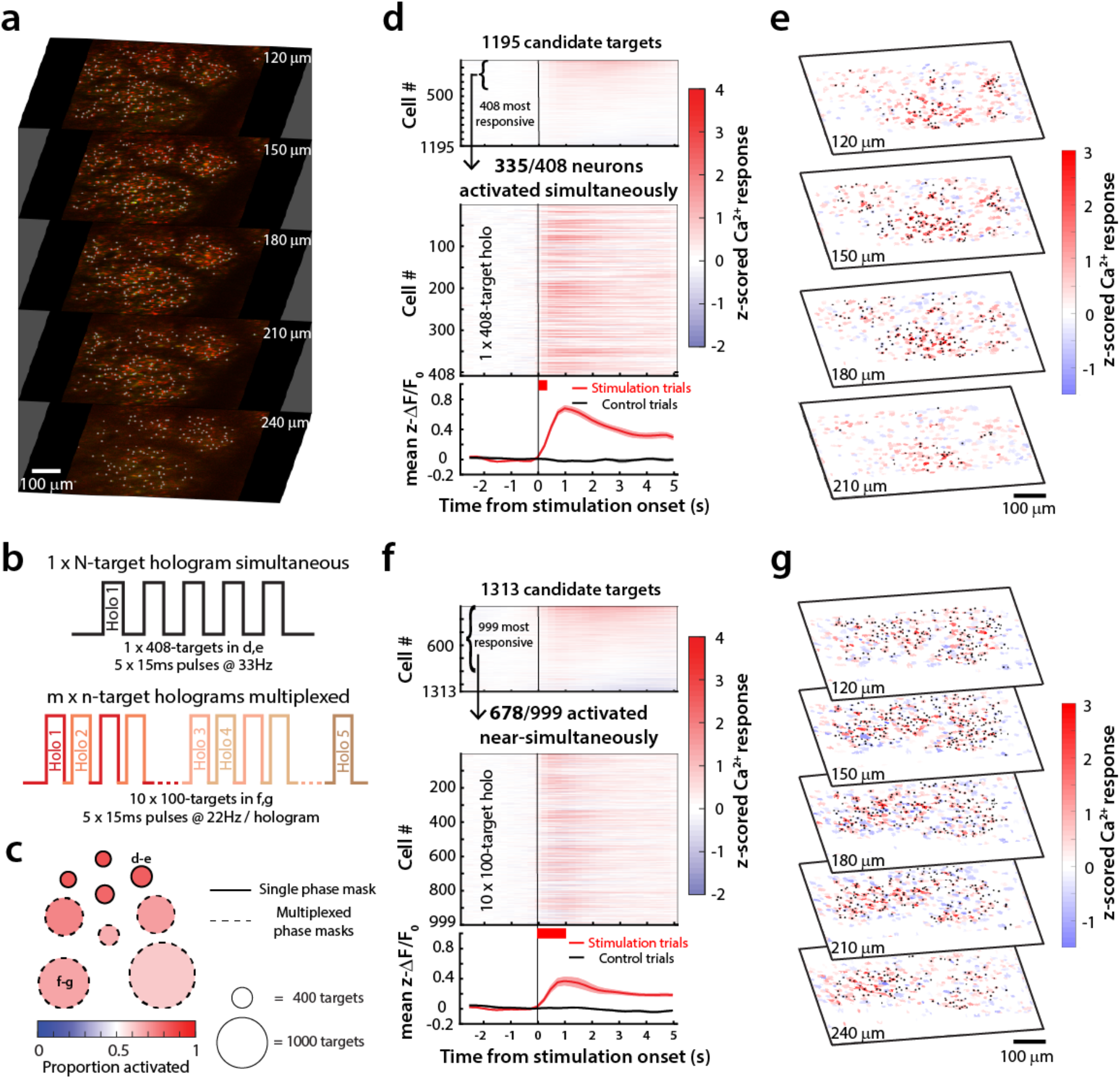
Large ensemble photo-stimulation in vivo with ChroME2s. **A)** Example 5-plane volume stack used for large ensemble photo-stimulation experiments in mice viral expressing ChroME2s-p2A-H2B-mRuby3. Initial targets for stimulation were identified based on nuclear mRuby3 expression (gray dots overlaid on red nuclei) and screened in small groups of 20-30 neurons for activation thresholds using 2-photon calcium imaging. **B)** Schematic of large ensemble stimulation protocols. Top: A single holographic ensemble comprised of the most activatable neurons in the volume was stimulated at a predetermined frequency (5 x 15 ms pulses at 33 Hz for the 408-target ensemble in **d**-**e**). Bottom: The most activatable neurons in the volume were randomly divided into *m* x *n*-target ensembles and stimulated in a multiplexed manner by fast switching of SLM frames (5 x 15 ms pulses at 22 Hz for the 10 x 100-target ensembles in **f**-**g**). **C)** Summary of the size and photoactivation of the targeted ensemble in each experiment using single phase masks (solid edged circles) and multiplexed phase masks (dashed edged circles). The diameter of the circle indicates the size of the ensemble, ranging from 308 targets for the single phase mask experiments to 1300 targets for the multiplexed experiments. The fill color indicates the proportion of targeted neurons successfully activated in each experiment. The example experiments shown in the remaining panels are denoted with the corresponding panel letters. **D)** Top: Raster plot of the response of 1195 candidate neurons across 4 planes to stimulation in the power screening step. In this example, the top 408 highly activated neurons at 20mW (Wilcoxon ranksum test between z-scored DF/F before and after stimulation, p < 0.05) were picked for subsequent large ensemble stimulation. Middle: Raster plot of calcium responses to the 408-target ensemble stimulation at 20mW. The thick red bar below the x-axis indicates the total duration of photostimulation. 335/408 neurons in the ensemble were significantly activated and 48/408 neurons were suppressed by the stimulation (Wilcoxon ranksum test, p < 0.05). Bottom: mean +/-s.e.m calcium response of all neurons in the targeted ensemble on stimulation trials (red trace) and control trials (black trace). **E)** 4-plane volumetric maps of the mean calcium responses of all recorded neurons for the 408-target ensemble experiment in **d**. **F)** Top: Raster plot of the response of 1313 candidate neurons across 5 planes to stimulation in the power screening step. In this example, the top 999 highly activated neurons at 45mW were picked for multiplexed large ensemble stimulation. Middle: Raster plot of calcium responses to the 10 x 100-target multiplexed ensemble stimulation at 45mW. The thick red bar below the x-axis indicates the total duration of photostimulation. 678/999 neurons in the ensemble were significantly activated and 219/999 neurons were suppressed by the stimulation (Wilcoxon ranksum test, p < 0.05). Bottom: mean +/-s.e.m calcium response of all neurons in the targeted ensemble on stimulation trials (red trace) and control trials (black trace). **G)** Similar to **e** but for the multiplexed ensemble stimulation shown in **f**.

In many cases, simply elevating firing rate across a large population of neurons in a single trial even without strict simultaneity (i.e., coactivation within a few ms) can be highly informative. We generated a series of holograms where each hologram targets ∼100 neurons and updated the SLM at a high rate that our simulations suggested could yield the maximum number of co-activatable neurons on a single one second epoch. Under these constraints, we were able to drive increases in firing rates in 667 neurons for 10 x 100-target holograms multiplexed into a single 1-second trial (Fig. 8f,g; Wilcoxon rank-sum test, one-tailed p < 0.025). Again, many of the targeted neurons showed net suppression during this large scale stimulation, presumably due to local, recurrent inhibition.

Taken together, the engineering and identification of microbial opsins optimized for patterned illumination optogenetics, combined with new paradigms for maximizing neural ensemble control, substantially expands the capabilities of spatially precise population neural control. Our data demonstrate that ChroME2f and ChroME2s are ultra-potent opsins that substantially increase the size of a controllable neural population with patterned illumination optogenetics, yet still provide outstanding millisecond temporal control across a broad array of conditions.

## Discussion

The biophysical properties of microbial opsins critically determine the scale, speed, and fidelity of optogenetic experiments, particularly for two-photon optogenetic excitation of neuronal population *in vivo*. Although numerous opsins have been characterized to date, there is still a need for opsins with enhanced features, specifically for multiphoton approaches where an ever-increasing scale and speed of photo-activation is desirable. We used rational design of the previously characterized opsin ChroME to design and validate a panel of mutants with enhanced properties, notably improved potency but with fast kinetics, and variants that span the spectral range, enabling all-optical experiments with the ever-expanding array of fluorescent optical indicators of neuronal activity, such as for calcium, voltage and neurotransmitter release, that span the two-photon spectrum. Importantly, we identified two ChroME variants, ‘ChroME2f’ and ‘ChroME2s’ that provide markedly enhanced photocurrents when expressed in cell culture or in cortical neurons *ex vivo*, yielding much greater optical sensitivity while preserving the temporal fidelity afforded by their parent opsin, ChroME. We present detailed two-photon excitation spectral analysis of these opsins, as well as provide the most comprehensive two-photon excitation spectral data on a set of other widely used opsins available to data. This work establises a critical knowledge base for a wide toolkit of opsins that can be used under a variety of experimental constraints.

In brain slices, ChroME2f and ChroME2s enabled very high-fidelity temporal control across a broad range of frequencies while requiring substantially less power than ChroME. Although ChRmine offers the ability to co-activate the most neurons with a limited power budget, it only provided reliable temporal control under a very narrow range of the conditions we tested, and primarily only with shorter illumination times that negate some of the benefit of its high sensitivity. Previous work has found that opsins with varying kinetic characteristics can enable sub-millisecond control under specific conditions(Chen *et al*., 2019, Ronzitti, Conti, *et al*., 2017, Marshel *et al*., 2019), although other work has demonstrated that under demanding applications opsins with faster closing kinetics more reliably provide high frequency, high fidelity temporal control(Gunaydin *et al*., 2010, Mager *et al*., 2018, Mardinly *et al*., 2018).

These studies, consistent with the work here, showed that opsins with fast closing kinetics limit extra spikes and missed spikes. Thus, when an experiment demands faithful reproduction of specific spike trains or frequencies of action potential generation, but when careful tuning of laser power or duration for each neuron is not possible, ChroME2f or ChroME2s are likely to be preferable to slower closing-kinetic opsins, such as ChRmine. Opsin expression levels can vary widely across neurons in a single preparation, and so tuning the light levels delivered to each neuron may be challenging. Among the opsins tested, ChroME2f appears to be most forgiving in that the reproduction of spike trains across frequencies did not appear power dependent (Fig. 3e). Conversely, when the specific aim is to activate as many neurons as possible at a time, ChRmine presents a potentially better choice. It is important to note that in this study we only used ‘wide-field’ large spot illumination of opsin-expressing neurons, rather than ‘spiral-scan’ activation. Whether these different modes of two photon optogenetic activation would yield different results may be tested in the future.

A key concern for any opsin in an all-optical experiment, however, is unwanted activation of the opsin by the imaging laser. Under identical conditions and among the four opsins we tested, our data show that ChRmine-expressing neurons exhibited the highest levels of such optical-crosstalk, with direct drive of action potentials depending on the imaging parameters. Although not as severe in all conditions, ChroME2s-expressing were also susceptible to substantial unwanted depolarization. In contrast, ChroME2f and ChroME exhibited similar levels of optical-cross talk, and substantially less than ChRmine or ChroMEs under most conditions. Under imaging parameters than can be used for large scale imaging of neuronal activity with calcium indicators - namely very large fields of view (∼1 mm x 1 mm), low frame rates (3-6 Hz), and modest imaging powers, unwanted depolarization for ChroME and ChroME2f was negligible, while for ChroME2s and ChRmine might still be deemed acceptable (∼1-5 mV). In contrast, when imaging smaller fields of view (∼0.3 mm 0.3 mm) at high frame rates (30 Hz), conditions that are also commonly used, the calcium imaging laser will likely directly drive action potentials in neurons expressing ChRmine and ChroME2s which may alter network physiology and thus compromise the interpretations of the experiment. These experiments argue that opsins and imaging conditions must be chosen with care to avoid unwanted sub-threshold, and occasionally supra-threshold, activation of opsin-expressing neurons by imaging lasers(Packer *et al*., 2014). Notably, we only tested such ‘optical cross-talk’ in L2/3 pyramidal cells. Other classes of neurons which may exhibit significantly higher intrinsic excitability (such as neurons with higher input resistance, lower action potential thresholds, or resulting from higher opsin-expression levels) might be activated under a broader array of conditions. Thus, it may be important for any given experimental paradigm to directly test this in the cell types studied and under the imaging conditions used.

*In vivo* we found that ChroME2f and ChroME2s substantially outperformed ChroME with respect to the power it took to photo-activate neurons. Correspondingly, we found that with ChroME2s we could simultaneously activate very large populations of neurons in the brains of awake mice. Brain heating places an upper limit on the scale of such experiments(Picot *et al*., 2018, Owen, Liu and Kreitzer, 2019, Mardinly *et al*., 2018, Podgorski and Ranganathan, 2016), and thus further engineering or identification of even more potent opsins could still be advantageous depending on the goals of an experiment. Another constraint in computer generated holography is that as the number of target spots increases in a hologram, the contrast ratio of the hologram decreases(Pégard *et al*., 2017). This can be addressed by using sparser holograms and rapidly multiplexing between phase masks with ultrafast SLMs, using multiple conventional SLMs(Marshel *et al*., 2019), or by scanning the excitation laser across a single SLM that is split along its length into separate phase masks(Parot *et al*., 2020).

Regardless, the high-performance opsins we present here provide the ability to co-activate much larger ensembles but still with very high temporal fidelity than previously possible. This opens the door to a much broader array of perturbations that should help elucidate the neural basis of perception, cognition and behavior.

## Author contributions

S.S. developed and tested all novel opsin constructs in cell culture and brain slices (Figure 1,2). H.A. performed power and temporal fidelity experiments (Figure 3,4) and opsin spectra characterization (Fig. 6). M.G and M.S. performed all optical cross-talk experiments (Figure 5). M.O. conducted the *in vivo* comparison of opsins (Figure 7). U.J. performed large-scale *in vivo* photo-stimulation experiments (Figure 8). L.A. designed and implemented various optical paths for opsin testing. I.O., H.B., W.H. and L.A. contributed to development of *in vivo* photo-stimulation parameters, wrote and tested software to operate the microscopes, pilot tested opsin constructs and expression parameters. I.T. generated and tested some opsin mutants. S.B. advised the design of mutant opsins. K.G. prepared mice via in utero electroporation and maintained the animal colony for all experiments. H.A. and all authors wrote and edited the manuscript.

## Acknowledgements

This work was funded by: NIH grant UF1NS107574 (HA), NIH Grant F32-EY031977 (WDH), the Simons Foundation Collaboration for the Global Brain award 415569 and NEI grant K99 EY029758-01 to I.A.O, and the New York Stem Cell Foundation. H.A. and S.B are New York Stem Cell Foundation Robertson Investigators. We thank Spectra Physics and Patrick Kolsch for use of the Insight X3, Karl Deisseroth for the ChRmine sequence and plasmid for cloning, and Janine Beyer for technical support. We thank all the members of the Adesnik lab for comments on the manuscript. The content is solely the responsibility of the authors and does not necessarily represent the official views of the National Institutes of Health.

---

**Figure S1:**
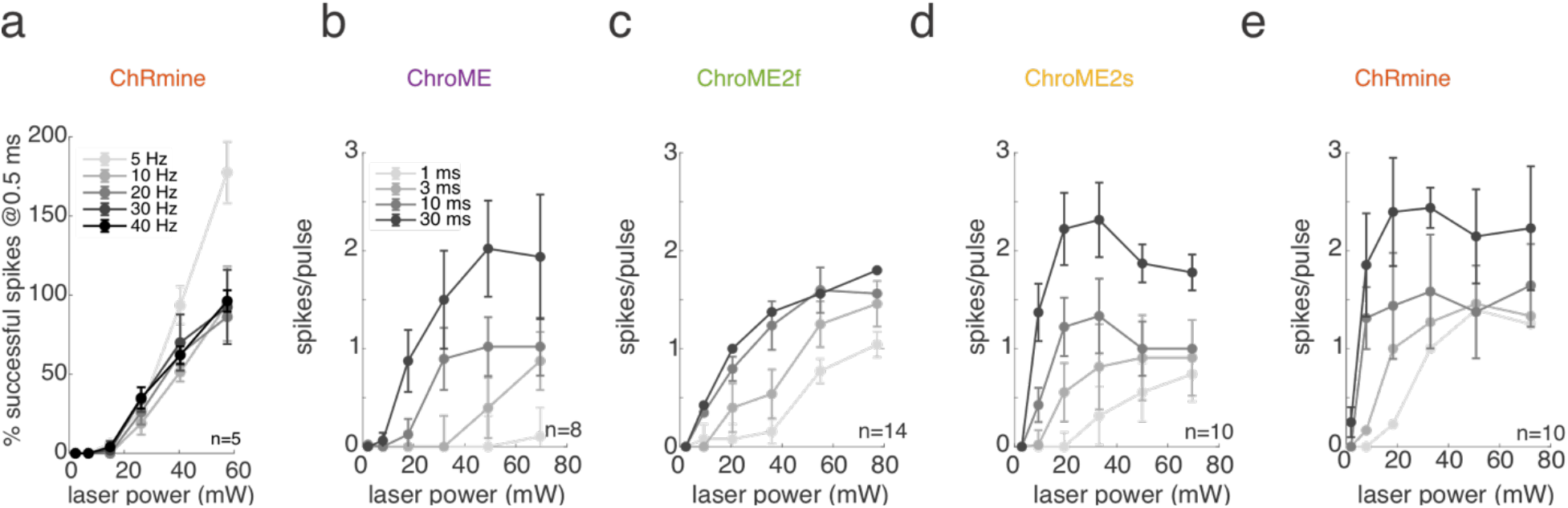
Additional characterization of two-photon-light evoked spiking in brain slices. a) The fraction of light pulses (duration = 0.5 ms) generating spikes across powers and stimulation frequencies for ChRmine-expressing cells when using fixed frequency pulse trains as in Figure 3. b-e) Light-evoked spikes per laser pulse as a function of laser power and pulse duration across the four indicated opsins.

**Figure S2:**
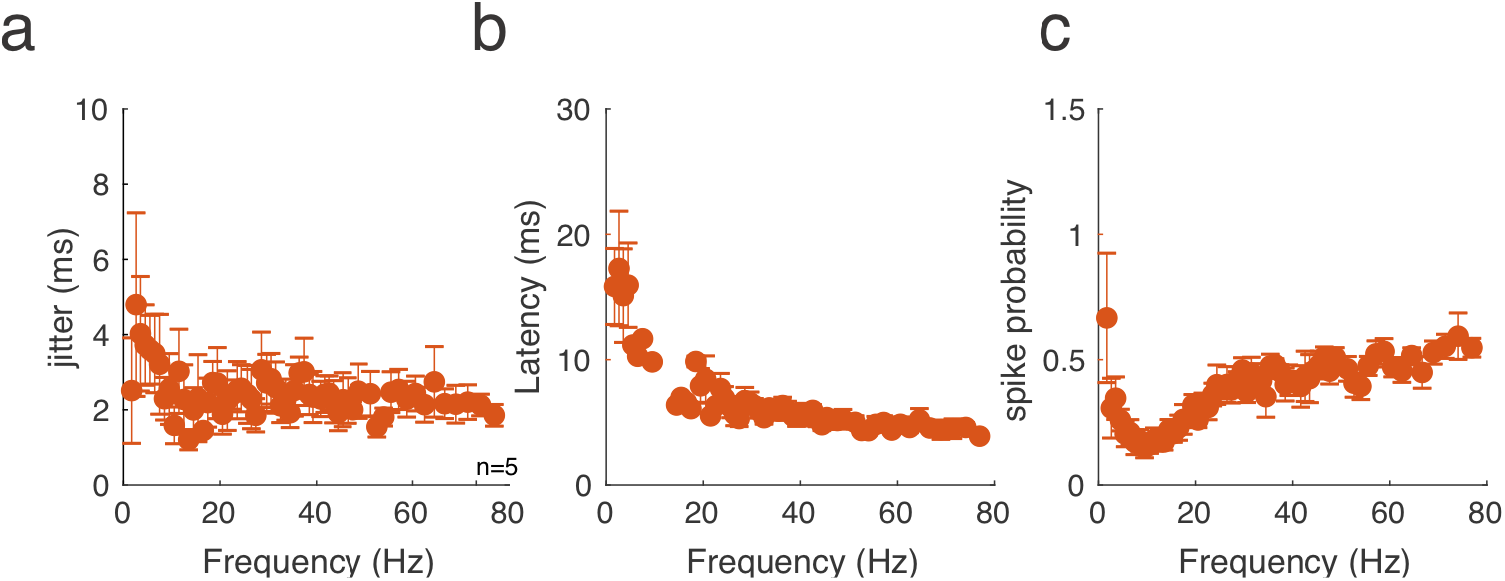
Frequency response of ChRmine-expressing neurons with 0.5 ms pulses of two-photon excitation. a) Jitter b) latency, and c) spike probability for light-evoked spikes (pulse duration = 0.5 ms) in ChRmine-expressing cells when using broadband poisson stimulation as in Figure 4.

**Figure S3:**
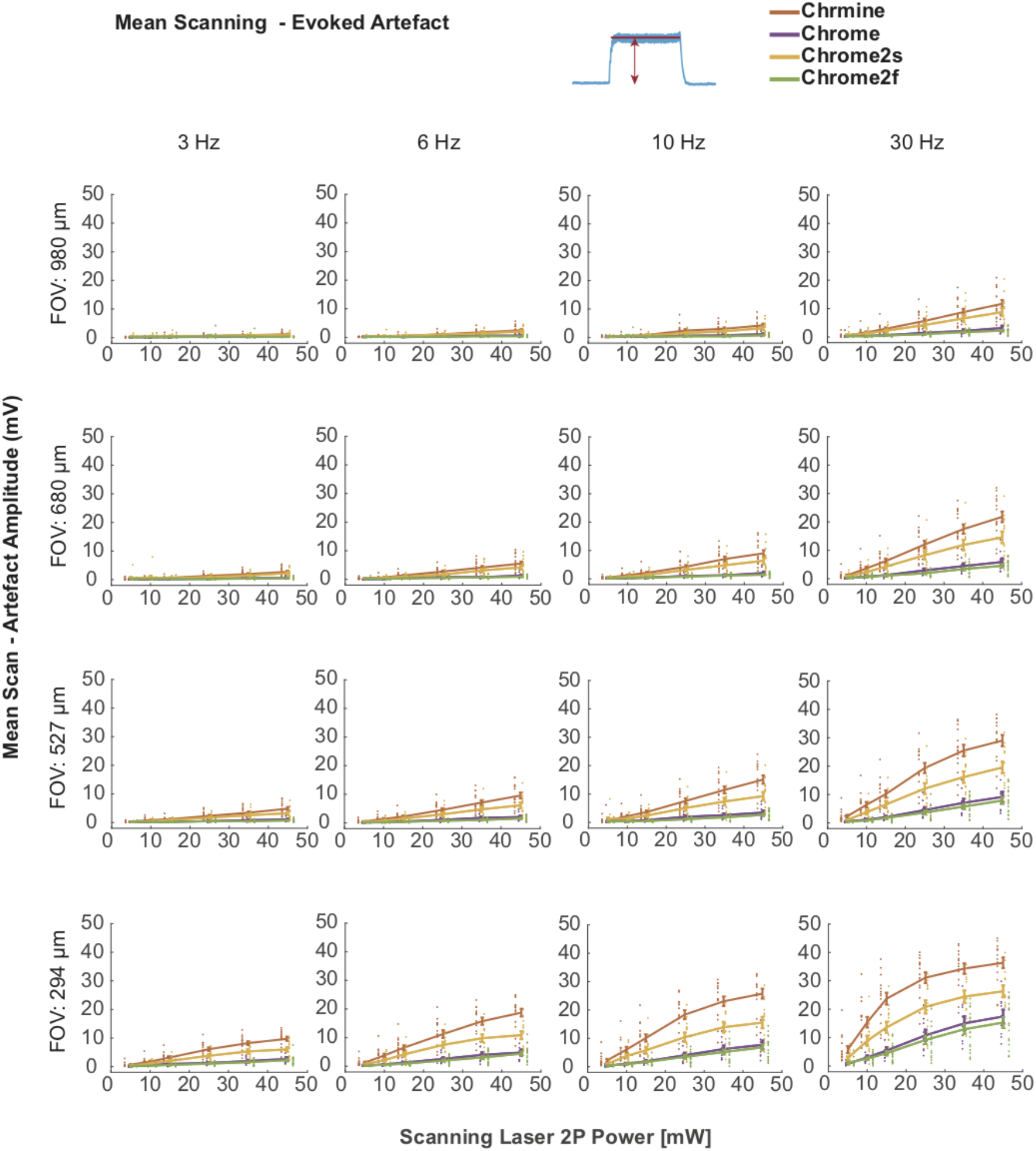
Mean raster-scanning evoked depolarization. Each plot shows the grand mean and standard error for all four tested opsins at various optical zooms and imaging frame rates. Dots specify the mean of single cells.

**Figure S4:**
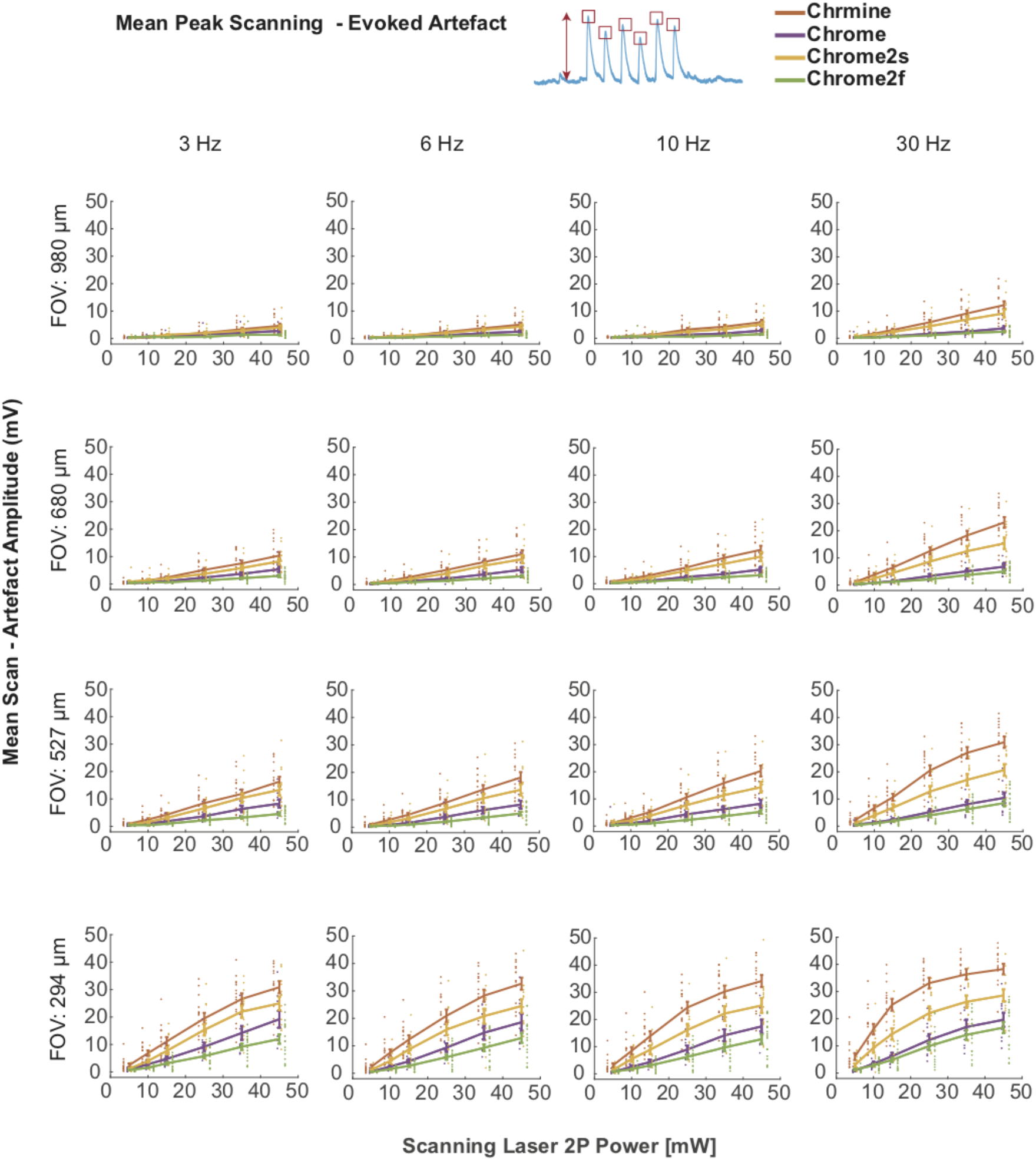
Average peak raster-scanning evoked depolarization. Each plot shows the grand mean and standard error for all four tested opsins at various optical zooms and imaging frame rates. Dots specify the mean of single cells.

**Figure S5:**
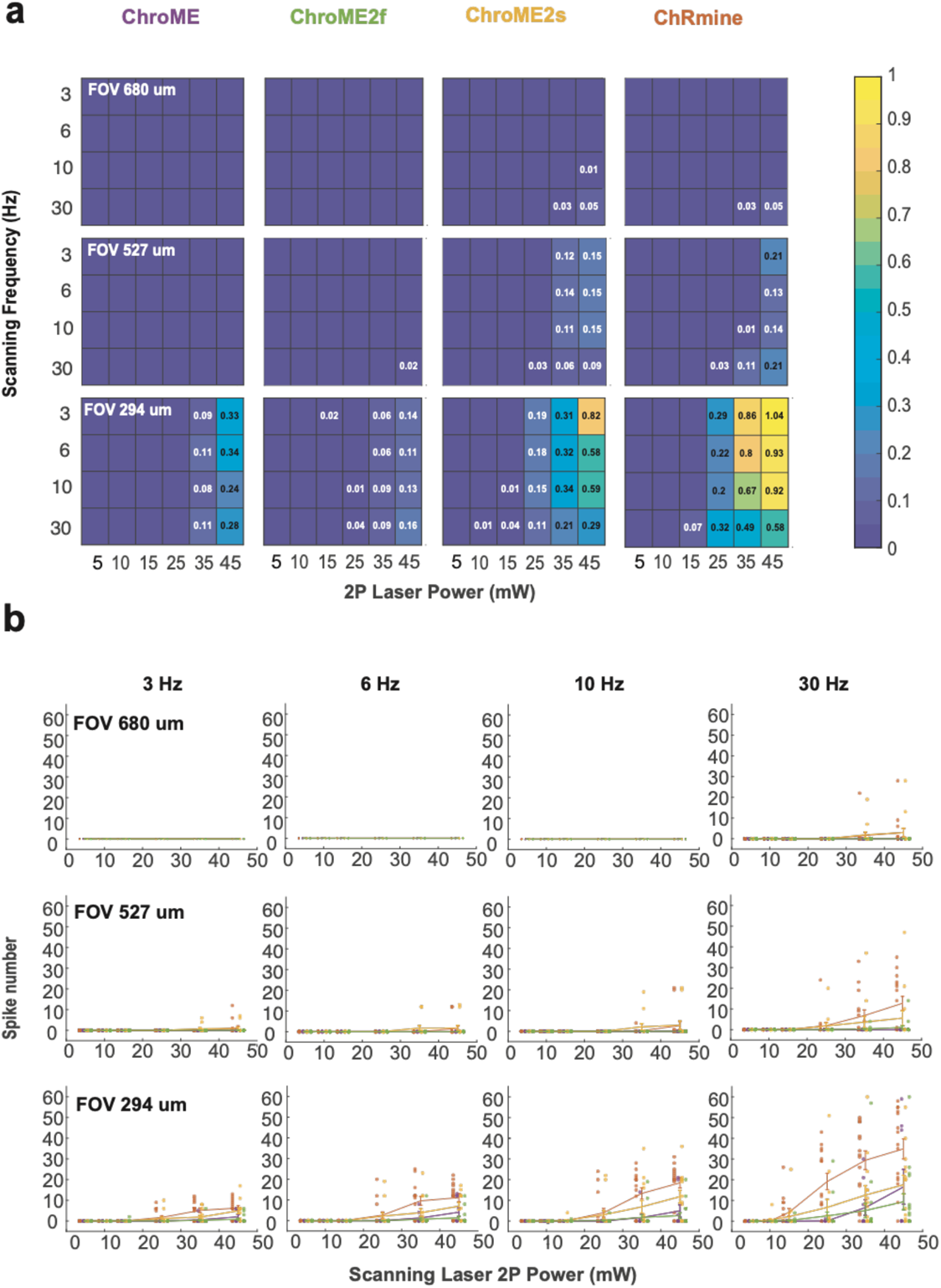
Raster-scanning evoked spiking across imaging conditions. A) Plots of the average number of spikes per frame-scan evoked at the three smaller FOVs for each opsin across scanning laser power. B) Same as in A) but with data from individual cells overlaid.

**Figure S6:**
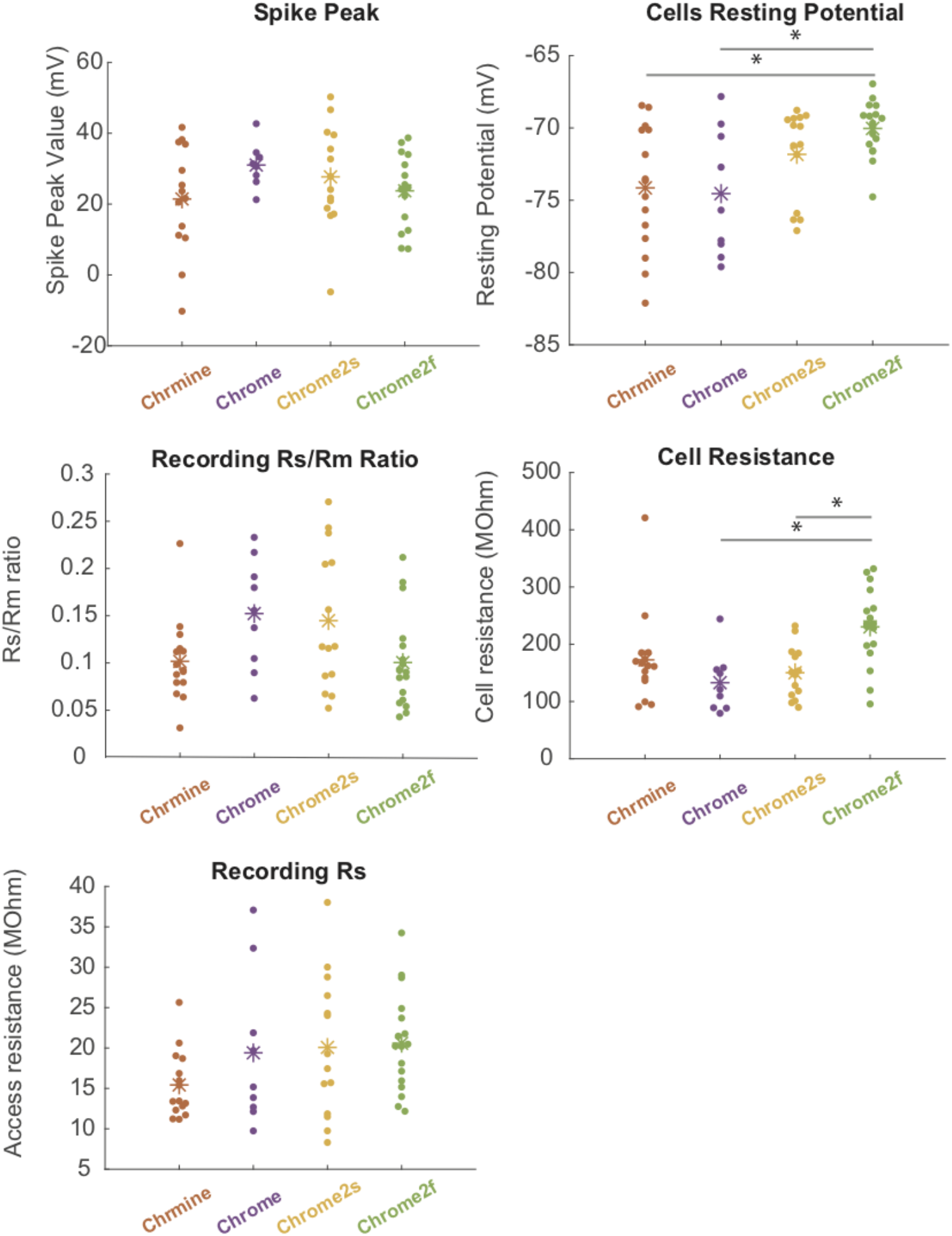
Physiological properties of the opsin-expressing L2/3 pyramidal cells. Peak spike voltage, mean resting potential, access/cell resistance ratio, mean cell membrane resistance, and access resistance for the neurons for which scanning evoked cross-talk was measured.

**Figure S7:**
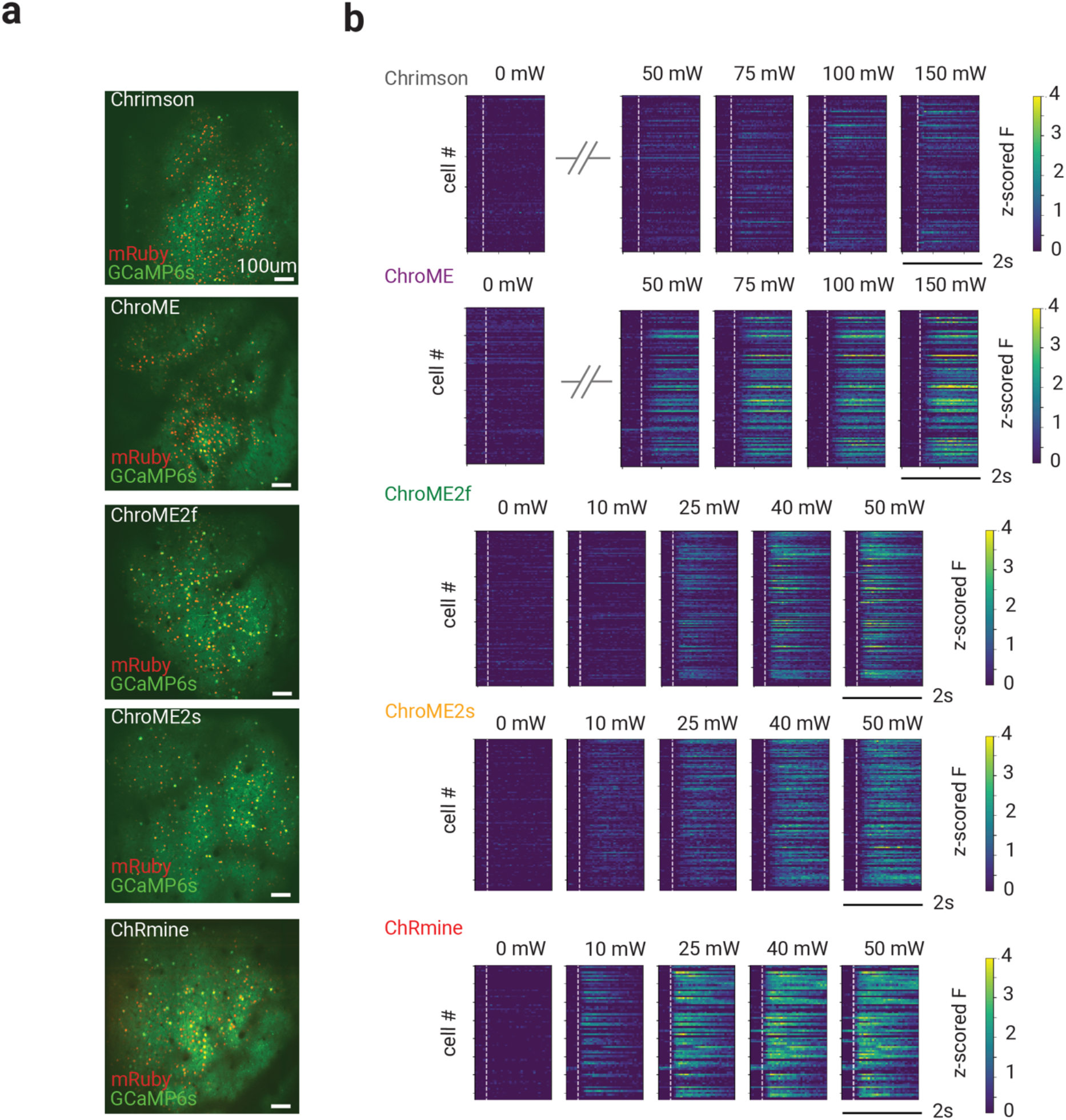
example images and calcium responses from all-optical opsin characterization experiments. **A)** *In vivo* two-photon images of a representative FOV from a mouse for each opsin, with excitatory neurons expressing GCaMP6s (green). Opsin expressing neurons are labeled with nuclear mRuby3 (red). Scale bar: 100mm. **B)** Representative mean z-scored fluorescence PSTHs for opsin-expressing cells stimulated at different powers. Dashed lines indicate the onset of photo-stimulation.

**Figure S8.**
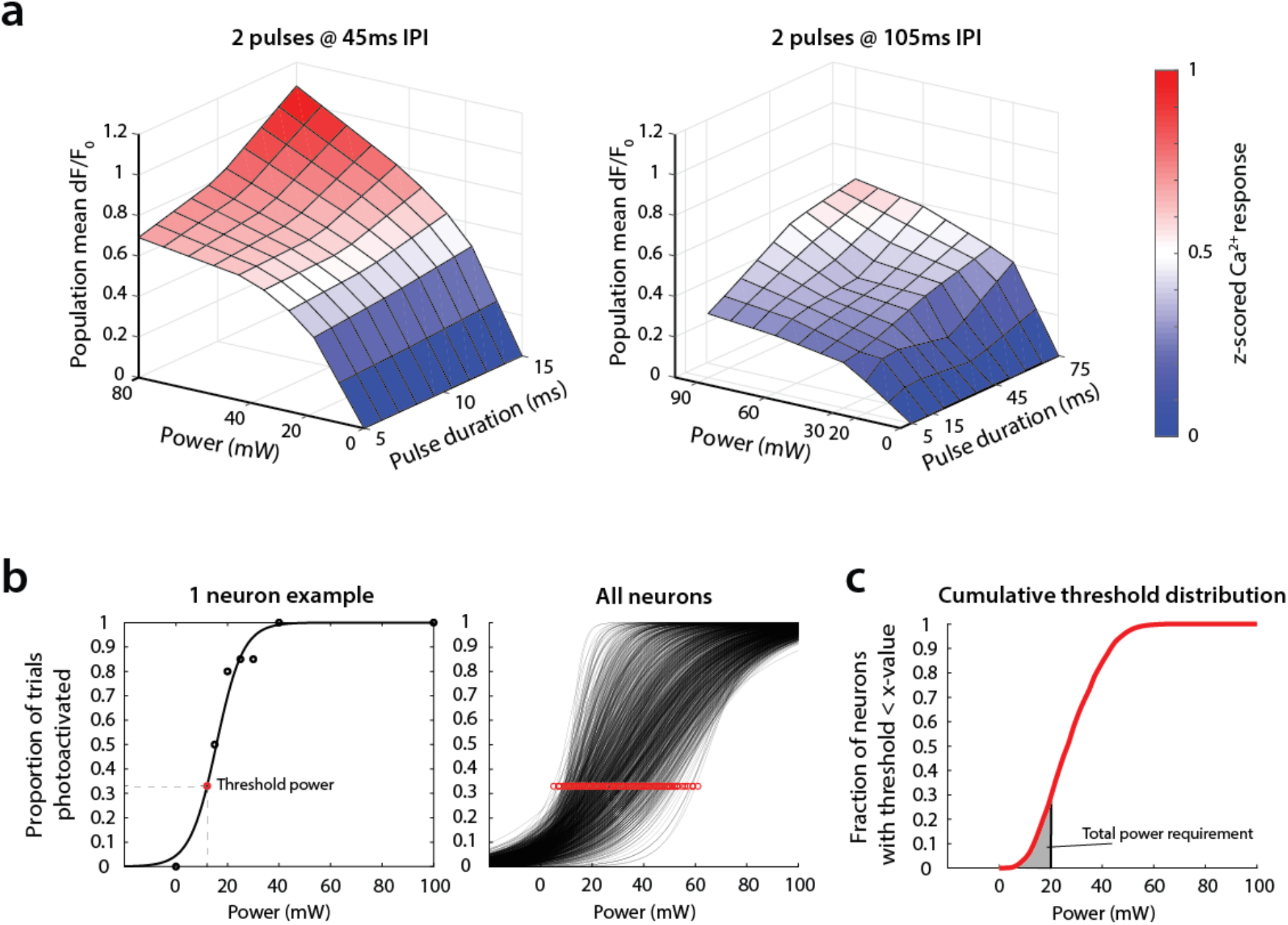
Stimulation parameters and power thresholds for large ensemble photo-stimulation experiments. **A)** Average parameter response surfaces used to determine optimal pulse width and frequency (parametrized as inter-pulse interval or IPI) for driving ChroME2s activation in large ensemble experiments. **B)** Prior to large ensemble stimulation, the optogenetic gain of each neuron in the volume was probed by targeting small groups at optimal pulse parameters determined above. Left: single neuron optogenetic response function depicting the proportion of trials in which the neuron was significantly photoactivated at each power (black circles – recorded proportions, black trace – logistic fit to circles the with an added anchor of 1 at 100mW). The red circle denotes the power at which the neuron is photoactivatable on 33% of the trials based on the fit, i.e., the threshold power. Right: Photo-activatability power curves (black traces) and threshold powers (red circles) for all neurons in an experiment. **C)** The cumulative distribution function of threshold powers in a session was used to determine the number of neurons selected for targeting in the subsequent large ensemble experiment. The integrated power in the shaded region corresponds to the total power requirement to successfully photoactivate all neurons with threshold powers lower than the vertical line, and this was matched to the available power to determine ensemble size.

## Methods

All experiments on animals were conducted with approval of the Animal Care and Use Committee of the University of California, Berkeley. In all experiments we attempted to use male and female mice equally. Mice were obtained from Charles River or bred in-house.

### Plasmid construction and Mutagenesis

Mutations in Chronos and ChroME were introduced by either site-directed mutagenesis or overlap extension PCR and verified by DNA sequencing. ChrimsonR, ChR2, CoChR, and ChRmine were obtained from Addgene. All opsins were fused to mRuby2 at their C-terminus and sub-cloned into the pCAGGS expression vector by In-Fusion cloning (Clontech, Mountain View, CA). In order to target the opsins to the soma and proximal dendrites of neurons, the sequence encoding the proximal restriction and clustering domain of the Kv2.1 voltage-gated potassium channel consisting of amino acids 536–600 (soma-targeting; ST) was codon optimized, synthesized (Integrated DNA Technologies, Coralville, IA) and inserted at the C-terminus of mRuby2 by In-Fusion cloning. For Adeno Associated virus (AAV) preparations, soma-targeted opsin cDNA was fused to a FLAG tag for immunofluorescence and a nuclear mRuby3 via a P2A self-cleaving peptide into pAAV and virus was prepared either at the Penn Vector Core or at the UC Berkeley Vision Core’s Gene Delivery Module.

### In Utero Electroporations and brain slice recording

Electroporations were performed on pregnant CD1 (ICR) mice (E15, Charles River). For each surgery, the mouse was initially anesthetized with 5% isoflurane and maintained with 2.5% isoflurane. The surgery was conducted on a heating pad to maintain body temperature, and warm sterile phosphate-buffered saline (PBS) was intermittently perfused over the pups throughout the procedure. A micropipette was used to inject ∼2 μl of recombinant DNA at a concentration of 2 μg/μl into the left ventricle of each embryo’s brain (typically DNA encoding opsins with GCaMP6s at a concentration of 2:1). Fast-green (Sigma-Aldrich) was used to visualize a successful injection. Using platinum-plated 5mm Tweezertrodes (BTX Harvard Apparatus) electrodes connected by a Y-connector to the negative pole, both sides of the embryo’s head were gently grabbed and a third electrode connected to the positive pole was placed slightly shifted below lambda to target the visual cortex and electroporated with 6 pulses at 30 V with a 1s delay using an Electro Square Porator (BTX Harvard Apparatus). After the procedure, the mouse was allowed to recover and come to term, and the delivered pups were screened for GCaMP6s expression and allowed to develop normally. Acute coronal slices were prepared and recorded from mice (ages p10–29) as described. Mice were screened with a handheld 300 mW 594 nm laser and filter goggles for expression after decapitation and before slicing. After slicing, recordings were made from the slices with strongest expression from the densest area as judged by fluorescence.

### In vitro electrophysiology

*In vitro* slice recordings were performed on 300um -thick coronal slices coming from P12 to P44 (opsin characterization experiments) or 4-6 week old animals (cross-talk experiments) in utero electroporated at E15.5 with plasmids containing one of the opsin: ChRmine, ChroME, ChroME2s or ChroME2f and GCaMP6s (2:1 proportion). Opsin positive cells in L2/3 were identified under 1P conditions. Whole-cell patch-clamp protocols were performed in ACSF perfusion solution (in mM: NaCL 119, NaHCO3 26, Glucose 20, KCl 2.5, CaCl 2.5, MgSO4 1.3, NaH2PO4 1.3) in temperature-controlled (33C) conditions.

Patch pipette (4-7 MOhm) were pulled from borosilicate glass filaments (Sutter Instruments) and filled with K-gluconate solution (in mM: 110 K-gluconate, 10, HEPES, 1 EGTA, 20 KCl, 2MgCl2, 2 Na2ATP, 0.25 Na3GTP,10 Phosphocreatine,295 mOsm,pH=7.45) mixed with 5uL of 50 uM Alexa hydrazide 488 dye (cross-talk experiments only, Thermo Fisher Scientific). Data was recorded at 20 kHz using 700b Multiclamp Axon Amplifier (Molecular Devices). The headstage with the electrode holder (G23 Instruments) were controlled by Motorized Micromanipulator (MP285A, Sutter Instruments). All data was acquired and analyzed with custom code written in Matlab using the National Instruments Data Acquisition Toolbox.

### Viral vectors (for in vivo)

Triple transgenic mice expressing GCaMP6s and Cre recombinase in excitatory neurons were obtained by crossing CaMKII-tTA to teto-GCaMP6s<u>43</u> ;Emx1 mice.

These mice were injected with adenoviral vectors expressing soma-targeted opsin and the red fluorophore mRuby3 in a Cre dependent fashion. All viruses had identical scaffold: AAV9-CAG.DIO.Opsin-FLAG-ST.P2A.H2B.mRuby3.WPRE.SV40 where “Opsin” is one of the five opsins shown in Fig. 7: Chrimson, ChroME, ChroME2f, ChroME2s or ChRmine. Final titer used for injections were: 1.73E12, 0.75E12, 2.8E12, 8.5E14, 5.85E13 viral genome/ml respectively.

Custom made viral preparations for Chrimson, ChroME and ChroME2s were generated by Addgene, whereas viral preparations for ChroME2f and ChRmine were generated by BerkeleyVision Science Core, Gene Delivery Module Facility.

### Surgery for in vivo experiments

AAV vectors were injected intracortically in V1 and cranial window surgeries were performed immediately after. Briefly, mice were anesthetized with isoflurane (2%) and administered 2 mg/kg of dexamethasone as an anti-inflammatory and 0.05 mg/kg buprenorphine as an analgesic. The scalp was removed, the fascia retracted, and the skull lightly etched. Following application of Vetbond (3M) to the skull surface, a custom stainless steel headplate was fixed to the skull with two dental cements: Metabond (C&B) followed by Ortho Jet (Lang). After the dental cement dried, a 3-mm diameter craniotomy over the left primary somatosensory cortex was drilled, and residual bleeding stopped with repeated wet–dry cycles using sterile artificial cerebrospinal fluid, gauze, and Gelfoam (Pfizer). A window plug consisting of two 3-mm diameter coverslips glued to the bottom of a single 5-mm diameter coverslip (using Norland Optical Adhesive #71) was placed over the craniotomy and sealed permanently using Ortho Jet (Lang). Animals were allowed to recover in a heated recovery cage before being returned to their home cage. Seven days after surgery, animals were habituated to head-fixation under a freely moving circular treadmill, *and in vivo* all-optical opsin potency estimations were done after 21 days post-surgery to allow for saturated opsin expression.

### One-photon and Two-photon Opsin Characterization experimental rig

For slice and cell culture experiments characterizing various opsins features (Figures 1-4) we employed a Scientifica slice scope equipped with Spectra X light engine for one-photon experiments (Lumencor), providing various light wavelengths and a Femtotrain 1040-5 (Spectra Physics, 10 MHz) for two-photon excitation. The laser beam was custom fitted to the scope resulting in a single focused two photon laser spot targeted at the cell. The spot size was ∼12.5 microns and generated through conventional achromatic lenses and a 40X olympus water immersion objective. Power was controlled by a Pockels Cell under computer control and gated by a laser shutter (Thorlabs). Photo-stimulus width was 5 ms unless otherwise indicated.

### CHO cell recording

Chinese hamster ovary CHO cells were transfected as above with a pCAG-opsin-mRuby2-Kv2.1 plasmid. 1-2 days after transfection coverslips with transfected cells were transferred to the brain slice rig described above. One-photon photostimulation of cells was performed at 510nm for Chronos, ChroME, and ChoME variants, and ChRmine at at a power of 0.45 mW using a Spectra X light engine (Lumencor), 470 nm for ChR2 and CoChR and 630nm for ChrimsonR. Currents were measured at a holding potential of −60 mV. The time to peak current was measured from average currents, and decay time constants were measured by fitting the traces from stimuli offset to a single exponential.

### Two-photon excitation spectra of opsins

Spectra collection experiments were performed on CHO cells prepared as mentioned above with an Insight X3 tunable laser. Power was controlled and carefully calibrated across the entire spectrum with the Pockels cell and set to be below saturation for all opsins. Power calibration was performed with a Thorlabs thermal power meter through the microscope objective to account for all optics coatings in the path and the power spectrum of the laser. Wavelength was controlled by the serial interface via Matlab.

### Brain slice characterization of opsin response characteristics

For the broadband ‘poisson’ photo-stimulation tests, photo-stimulus width was 5 ms unless otherwise indicated, and on each trial and random pulse train (pulse times drawn from a poisson distribution with mean rates of 10, 20 and 30 Hz) were delivered to each neuron. At the beginning of the experiment the power of a single 5 ms (or 0.5 ms in Figure S1a, Figure S2) laser exposure was set to reliably drive across successive trials (inter-trial interval = 2.3 seconds).

Spike jitter was computed as the standard deviation of spikes times across all light-evoked spiked in the experiment. Spike probability was computed as the fraction of light pulses that drove one or more spikes. Spike latency was the time between the onsite of the laser exposure and the peak of the subsequent action potential prior to any subsequent laser exposure.

### Two-photon holographic setups

In vivo opsin characterization and large-scale photostimulation experiments were performed on two 3D-SHOT multiphoton holographic setups (refer to (Mardinly *et al*., 2018)for detailed description). The setups were both custom built around a commercial Sutter MOM microscope platform (Sutter Instruments) and combined a 3D photostimulation path, a fast resonant-galvo two-photon raster scanning imaging path and a wide-field one-photon epifluorescence/IR transmitted light imaging path. The stimulation and imaging beams were merged together using a polarizing beamsplitter placed before the microscope tube lens.

Femtosecond fiber lasers were used for photostimulation: Satsuma HP2 (1030nm, 2MHz, 350fs, Amplitude Systemes) for in vivo opsin characterization (setup 1) and Monaco 1035-80-60 (1040nm, 1MHz, 300fs, Coherent) for large scale photostimulation (setup 2). On both setups, the stimulation laser was directed onto a blazed diffraction grating (600l/mm, 1000nm blaze, Edmund Optics 49-570 or 33010FL01-520R Newport Corporation) for temporal focusing. In order to be able to utilize the total available laser power on setup 2 (60W laser output), the beam was enlarged by a 2.5 factor to prevent photodamage of the grating surface. The spot on the grating was relayed onto a rotating diffuser where it formed a temporally focused spot. The rotating diffuser was used to both randomize the phase pattern simprinted on the temporally focused spot and to expand the beam on the direction orthogonal to the temporal focusing direction and fully fill the spatial light modulator (HSP1920 192×1152 pixels Medowlark Optics).The SLM plane was relayed through 4f systems to the back aperture of an Olympus 20x water immersion objective, resulting in custom 2D or 3D distribution of temporally-focused spots at the focus of the objective. Holographic phase masks were calculated using the iterative Gerchber-Saxton algorithm(Gerchberg and Saxton, 1972) and intensity distribution was corrected to accommodate for diffraction efficiencies.

The two-photon imaging paths relied on Ti:sapphire lasers, Chameleon (Coherent, setup 1) or Mai Tai (Spectra Physics, setup 2), with external power control via Pockels cells (Conoptics, Inc). For fast raster scanning, both systems were equipped with conjugated 8kHz resonant galvo-galvo systems (relayed with either a pair of Plössl lenses -setup1- or a pair of 90° off-axis parabolic mirrors, setup2, Negrean and Mansvelder, 2014). The imaging path hardware was controlled by ScanImage software and custom Matlab code was used to control the spatial light modulator for targeted photostimulation and synchronize with imaging.

Epifluorescence excitation was via an X-Cite LED (Excelitas Technologies, patch clamp rig for Fig. 6) or Spectra X (Lumencor, patch clamp fig for Fig. 1-5) light source filtered by appropriate excitation filter set. For slice transillumination we used a 750nm and IR diffuser. The image was collected using an Olympus 20 × magnification water-immersion objective and a CCD camera and displayed on a screen enabling targeted patch clamping (Fig. 6) or Olympus 40x water immersion objective (Figs. 1-5).

### Cross-talk characterization data collection

Cross-talk characterization was performed in current clamp across 96 different imaging conditions of variable FOV (980, 680, 527,294 um), imaging frequency (3,6,10,30 Hz) and power (5,10,15,25,35,45 mW). Each experimental trace contained a baseline period and 2s scanning period at a given condition. At the beginning of the experiment, we used current injection to characterize each cell’s intrinsic properties. For voltage clamp experiments cells were clamped at -70 mV. Access resistance was monitored throughout the protocol’s duration and only cells in which the Rs/Rm ratio was below 0.3 were accepted (mean and standard error for each opsin group: ChRmine 0.1+/-0.01, ChroME 0.15+/-0.02, ChroME2s 0.15+/-0.02, ChroME2f 0.10+/-0.01). For current clamp protocols, cells were kept at resting potential and only cells with stable resting potential (<4 mV change) during the entire protocol were accepted for data analysis.

Each opsin group was collected using tissue obtained from 4-7 animals and contained: 15,9,14,17 cells for ChRmine, ChroME, ChroME2s, ChroME2f respectively. For all the measurements, data were baseline-subtracted and spikes were removed by applying a median filter. Outliers analysis was applied to the 2P evoked photocurrent results to exclude cells whose mean was 2 standard deviations away from the population mean at each power. The same group of cells was taken into account in all the measurements unless specified differently. All measurements are expressed as the grand mean over all the cells and standard error. One-way ANOVA across all conditions was calculated to specify statistical significance and post hoc analysis (Fisher’s Least Significant Difference test). For scanning-evoked spike count analysis scanning period was binned into 5 ms periods. Spikes were detected by crossings above 0 mV.

### In vivo all optical experiments

Mice were head-fixed on a freely spinning running wheel. Imaging and photostimulation wavelengths were respectively 920nm and 1030-1040nm. Neurons that co-expressed opsin and nuclear-targeted H2B-mRuby3 and GCaMP6s were automatically segmented based on frame-averaged images taken at 1020 nm using custom peak-intensity detection Matlab software. The centroids of the segmented masks were then used to compute holographic phase masks that were loaded sequentially on the spatial light modulator. For experiments in Fig. 7, acquisitions were performed at 15 or 30Hz frame rate (15Hz for ChroME2f, ChroME2s and ChRmine, 30Hz for ChroME and Chrimson) and with ∼900 x 900 µm fields of view. The imaging/photostimulation plane was between 100 and 200 µm below the pial surface and the imaging power was below 50mW for all opsins. Large ensemble stimulation experiments were performed on setup 2 (see description above). For these experiments, fast multi-plane imaging was achieved with an electrically tunable lens (Optotune AG) in the imaging path just before the scanning unit. Typically, 3-5 planes spaced 30µm apart were acquired at 4-5 Hz frame rate. Each plane covered a ∼700 x 700 um field of view, and the imaged volume had an axial span of 60-120um and was located between 100-300um below the pial surface. Stimulation targets were identified across all planes using the procedure described above.

### 2P imaging and photo-stimulation data analysis

#### Opsin comparison experiments

For power curve comparison experiments in Figure 7, motion correction, calcium source extraction, and deconvolution were performed using Suite2p(Pachitariu *et al*., 2016). Briefly, raw calcium videos were motion-corrected using Suite2P with subpixel alignment = 10, and calcium sources were extracted with key parameters diameter = 7–10um. Calcium sources were then manually examined and accepted or rejected based on their overlap with morphologically identifiable neurons. Neuropil subtracted fluorescence vectors (*F*) (using a neuropil coefficient of 0.7) were used for downstream analysis. Calcium signals were acquired continuously, and each cell’s fluorescence was *z*-scored. Holographic targets were aligned to calcium sources by calculating the Euclidean distance between the centroids of all holographic targets and all calcium sources and finding the minimum. Rarely, the automated software assigned targets to calcium sources with distance >10 µm; these were excluded from subsequent analysis. For power curve comparison experiments in Fig. 7, Neurons were stimulated in groups of 1-10 targets per phasemask. The stimulation consisted in 5 pulses of 5ms duration at 30Hz for each hologram, and all cells were stimulated at different powers.

### Large ensemble stimulation experiments

The mean z-scored F of each cell was aligned with the time of the stimulation, a baseline of 5-10 frames was calculated and compared to the mean z-scored F in the response window. A trial-wise two-sided Wilcoxon test was used to assess whether each cell was photoactivated.

The fraction of photoactivated cells was calculated by dividing successfully activated neurons by the total number of targeted neurons.

The simultaneous activation of many neurons introduced large and systematic correlations that led to a miscount of calcium sources based on Suite2P alone. Therefore, we employed a hybrid calcium source extraction procedure that combined Suite2P-based extraction as described above for the opsin comparison experiments (diameter = 14um, neuropil/cell diameter ratio = 5) with custom extraction of calcium signals around all previously identified holographic targets using the same parameters for cell and neuropil size that were used for the Suite2P step.

## Notes

### Competing Interest Statement

The authors have declared no competing interest.

